# Cytoplasmic CPSF6 regulates HIV-1 capsid trafficking and infection in a cyclophilin A-dependent manner

**DOI:** 10.1101/2020.06.05.136697

**Authors:** Zhou Zhong, Jiying Ning, Emerson A. Boggs, Sooin Jang, Callen Wallace, Cheryl Telmer, Marcel P. Bruchez, Jinwoo Ahn, Alan N. Engelman, Peijun Zhang, Simon C. Watkins, Zandrea Ambrose

## Abstract

Human immunodeficiency virus type 1 (HIV-1) capsid binds host proteins during infection, including cleavage and polyadenylation specificity factor 6 (CPSF6) and cyclophilin A (CypA). We observe that HIV-1 infection induces higher-order CPSF6 formation and capsid-CPSF6 complexes co-traffic on microtubules. CPSF6-capsid complex trafficking is impacted by capsid alterations that reduce CPSF6 binding or by excess cytoplasmic CPSF6 expression, both of which are associated with decreased HIV-1 infection. Higher-order CPSF6 complexes bind and disrupt HIV-1 capsid assemblies *in vitro*. Disruption of HIV-1 capsid binding to CypA leads to increased CPSF6 binding and altered capsid trafficking, resulting in reduced infectivity. Our data reveal an interplay between CPSF6 and CypA that is important for cytoplasmic capsid trafficking and HIV-1 infection. We propose that CypA prevents HIV-1 capsid from prematurely engaging cytoplasmic CPSF6 and that differences in CypA cellular localization and innate immunity may explain cell-specific variations in HIV-1 capsid trafficking and uncoating.

**Graphical Abstract:** 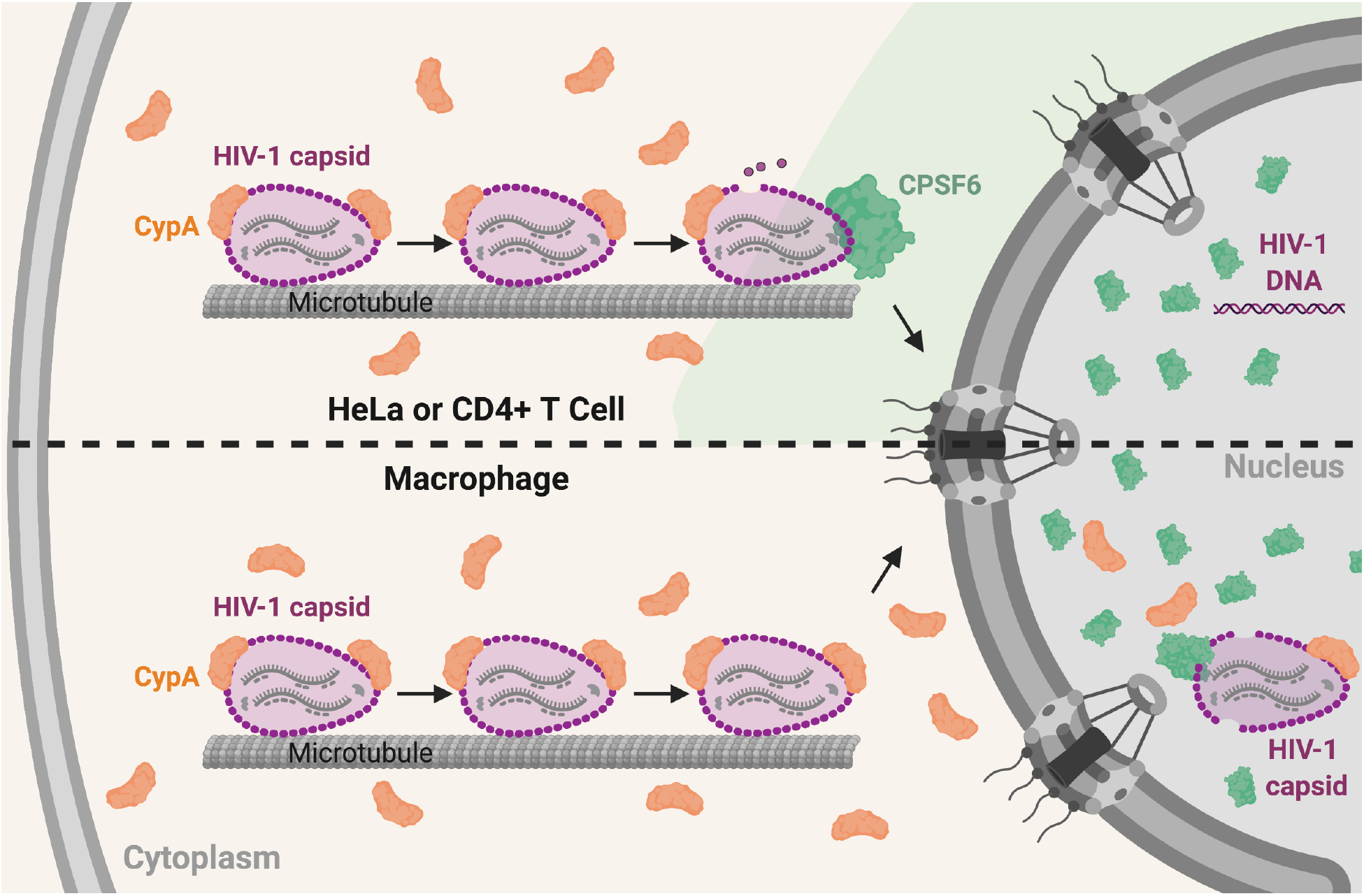

## Introduction

Human immunodeficiency virus type 1 (HIV-1) capsid is a unique structure that assembles after viral proteolytic cleavage of the Gag and Gag-Pol polyproteins during or after virus budding and release from cells. Multiple HIV-1 capsid proteins (CAs) form a conical-shaped core consisting of approximately 200 hexamers and 12 pentamers, which encapsulates two copies of the RNA genome (Ganser et al., 1999; Zhao et al., 2013). An important yet understudied aspect of HIV-1 infection, which is referred to as uncoating, describes the process of capsid dissociation that occurs during reverse transcription and before viral DNA integration into host chromatin (Ambrose and Aiken, 2014; Campbell and Hope, 2015). HIV-1 capsid uncoating is dependent on microtubule trafficking (Lukic et al., 2014; Malikov and Naghavi, 2017; Pawlica and Berthoux, 2014) and may occur in a multi-step process (Xu et al., 2013). HIV-1 infection requires active transport of the viral reverse transcription complex/preintegration complex (RTC/PIC) through the cellular nuclear pore complex (NPC) (Bukrinsky et al., 1992), and the majority of CA is likely uncoated at the NPC and/or within the nucleus (Arhel et al., 2007; Burdick et al., 2017; Burdick et al., 2020; Francis and Melikyan, 2018; Zurnic Bonisch et al., 2020). Single mutations in CA can greatly impact the stability of HIV-1 capsid, altering its uncoating and affecting virus infectivity (Rihn et al., 2013; von Schwedler et al., 2003).

HIV-1 capsid has been shown to bind several host proteins during infection (Yamashita and Engelman, 2017). Two examples are cyclophilin A (CypA) (Gamble et al., 1996; Luban et al., 1993), which is relatively abundant in cells (Koletsky et al., 1986), and cleavage and polyadenylation specificity factor 6 (CPSF6) (Lee et al., 2010). Disruption of HIV-1 capsid binding to CypA can occur via amino acid substitution, such as G89V and P90A, in the loop between helices 4 and 5 in CA (Franke et al., 1994; Wiegers et al., 1999) or by treatment with small molecule inhibitors, such as cyclosporine A (CsA) (Luban et al., 1993). Inability to bind to CypA in target cells can affect the infectivity of the invading HIV-1 particle (Sokolskaja et al., 2004). CypA dependence is cell type specific and has been shown to affect multiple steps in the virus life cycle, including capsid uncoating, reverse transcription, nuclear import, or integration (Braaten et al., 1996; De Iaco and Luban, 2014; Qi et al., 2008; Schaller et al., 2011).

CPSF6 is an arginine/serine (RS)-domain containing protein expressed predominantly in the cell nucleus and is involved in splicing and polyadenylation of host RNAs (Dettwiler et al., 2004). CPSF6 binds to HIV-1 capsid at an interface between two CA monomers defined by helices 3, 4, and 5 (Lee et al., 2010; Price et al., 2012). The beta-karyopherin transportin 3 (TNPO3) is the key mediator of RS domain protein nucleocytoplasmic transport in cells, and TNPO3 directly binds the C-terminal RS domain in CPSF6 to affect its nuclear import (Jang et al., 2019; Maertens et al., 2014). Accordingly, truncation of the C-terminus of CPSF6 leads to increased cytoplasmic expression and inhibition of HIV-1 nuclear entry and infection (Jang et al., 2019; Lee et al., 2010) in a TNPO3-dependent manner (De Iaco et al., 2013; Fricke et al., 2013). Disruption of CPSF6 binding to HIV-1 capsid is mediated by alteration of CPSF6 at amino acid F321 in the central proline-rich domain (Price et al., 2014) or by mutations in CA, including residues 57, 70, 74, 77, and 105 (Henning et al., 2014; Lee et al., 2010; Price et al., 2012; Saito et al., 2016). While reduction of CPSF6 expression or loss of capsid binding to CPSF6 in cells does not affect overall HIV-1 infection in most cell types, viral DNA integration is mistargeted outside of gene dense, transcriptionally active host chromatin to heterochromatic lamina-associated domains (Achuthan et al., 2018; Schaller et al., 2011; Sowd et al., 2016).

In this study, we investigated whether HIV-1 capsid interacts with CPSF6 in the host cell cytoplasm and whether this interaction affects capsid trafficking and subsequent virus infectivity in different cell types. Using live-cell microscopy, we visualized wild type (WT) HIV-1 complexes colocalized with cytoplasmic CPSF6 that trafficked together on microtubules. By negative-stain transmission electron microscopy (TEM), we show that purified CPSF6 protein forms oligomers that bind and disrupt CA tubular assemblies. Inhibiting HIV-1 capsid interaction with CypA led to increased association of viral particles or *in vitro* CA assemblies with CPSF6 and changes in WT HIV-1 complex trafficking that corresponded to reduced infectivity. Depletion of CPSF6 affected capsid trafficking, albeit differentially depending on the cell type.

## Results

### CPSF6 is expressed in the perinuclear region and traffics on microtubules with WT HIV-1 complexes

As CA protein may dissociate from HIV-1 nucleic acid complexes prior to entry into the nucleus where CPSF6 predominantly is expressed, we examined whether CPSF6 was expressed in the cell cytoplasm. Antibody staining of endogenous CPSF6 (NBP1-85676, Novus) or expression of green fluorescent protein (GFP)-tagged CPSF6 (CPSF6-GFP) in HeLa cells showed mostly nuclear expression as well as punctate cytoplasmic expression mainly near the nuclear membrane, which may indicate higher-order complex formation (**Fig 1A**). Highly Inclined and Laminated Optical sheet (HILO) live-cell microscopy showed that perinuclear CPSF6-GFP puncta were dynamic in cells and were co-localized with microtubules (**Fig 1B, Movie S1**). Inhibition of microtubule polymerization with nocodazole inhibited CPSF6-GFP movement in cells, suggesting that CPSF6 traffics on microtubules (**Fig 1C**).

**Figure 1.**
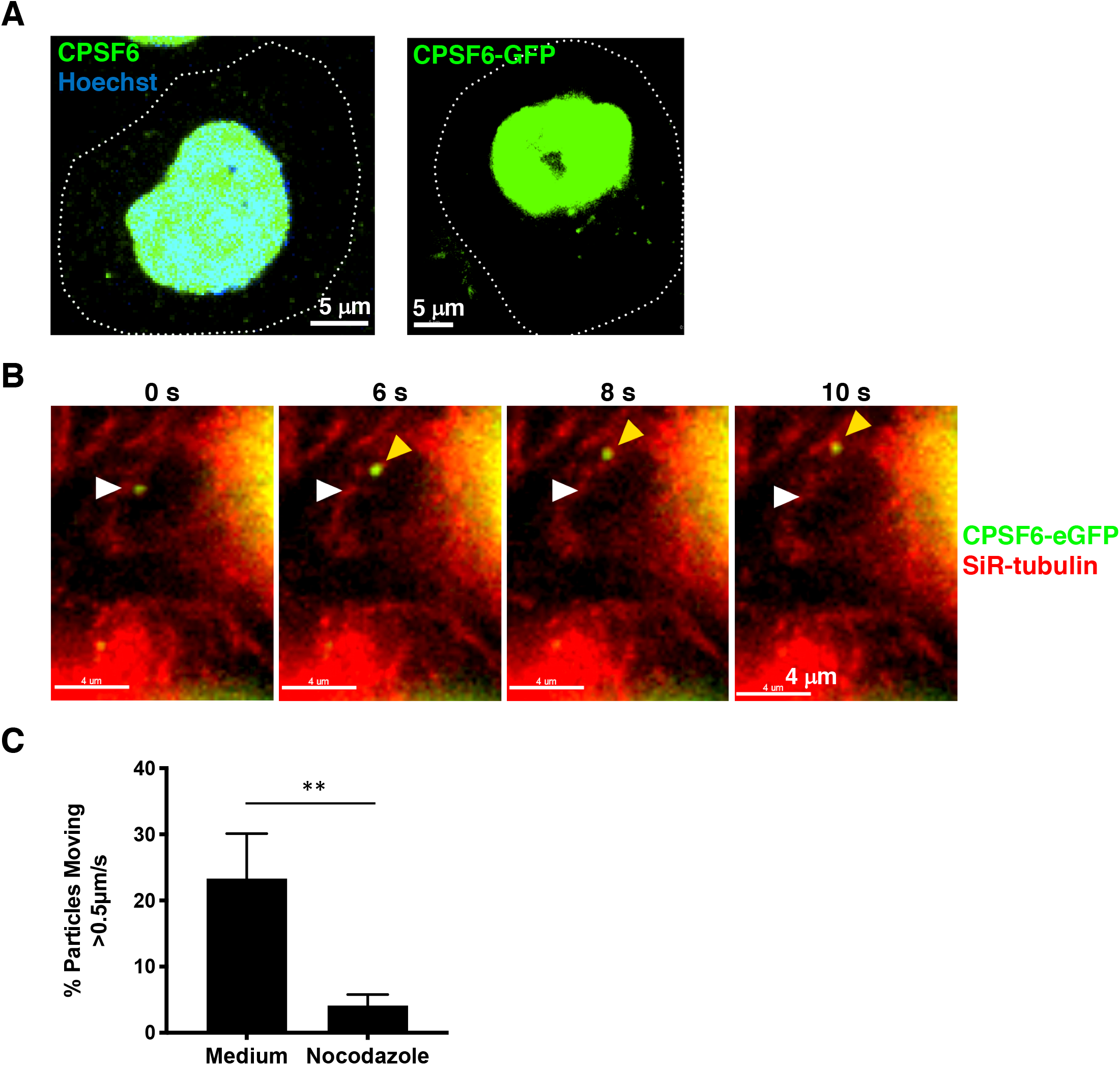
CPSF6 puncta are detected in the perinuclear region and traffic on microtubules. (A) Endogenous CPSF6 or expression of CPSF6-GFP in HeLa cells (dotted lines, cell outlines). CPSF6 is expressed as two different isoforms composed of 551 or 588 amino acid residues (Ruegsegger et al., 1998); exogenously expressed proteins throughout this study were based on the 588 isoform. (B) Movement of a CPSF6-GFP higher order complex (green) is shown in a HeLa cell stained with SiR-tubulin (red) by HILO live-cell imaging. The white arrow indicates the location of the complex at the first time point and yellow arrow indicates the location at subsequent time points. (See also Movie S1.) (C) The percentage of cytoplasmic CPSF6-GFP complexes are shown that trafficked > 0.5 μm/s in HeLa cells treated with or without nocodazole. Error bars indicate standard deviations (STDEV) of n ≥ 185 complexes.

As HIV-1 RTCs also traffic on microtubules (Dharan and Campbell, 2018) (**Movie S2**), we examined the association of fluorescently labeled HIV-1 particles with cytoplasmic CPSF6-GFP in cells. Vesicular stomatitis virus glycoprotein (VSV-G)-pseudotyped WT or N74D HIV-1 encoding firefly luciferase and labeled with integrase (IN) tagged with the fluorophore mRuby3 or tagRFP was used to infect cells expressing CPSF6-GFP. Labeled IN co-localized with viral RNA in particles or with viral RNA and CA protein early after infection, the latter of which demarcates RTCs/PICs (Ning et al., 2018). WT complexes were co-localized with perinuclear CPSF6-GFP, while N74D complexes were not (**Fig 2A**). Multiple WT or N74D virus particles that co-localized with CPSF6-GFP were assessed by live-cell imaging over time. The fluorescence intensity of WT HIV-1 complexes remained associated with CPSF6-GFP, whereas the fluorescence intensity of N74D viral particles did not (**Fig 2B**). This is consistent with a recent study showing WT HIV-1 particles associated with CPSF6 initially outside of the nucleus (Burdick et al., 2020). WT HIV-1 particles associated with CPSF6-GFP trafficked rapidly and linearly, suggestive of microtubule movement (**Fig 2C, Movie S3**). Consistent with this interpretation, trafficking of WT HIV-1 particles associated with CPSF6-GFP was inhibited with nocodazole treatment (**Fig 2D**). Finally, CPSF6 tagged with iRFP670 trafficked towards the nucleus with its TNPO3 binding partner, which was tagged with GFP (**Fig 2E, Movie S4**). These results suggest that HIV-1 complexes traffic with CPSF6 on microtubules.

**Figure 2.**
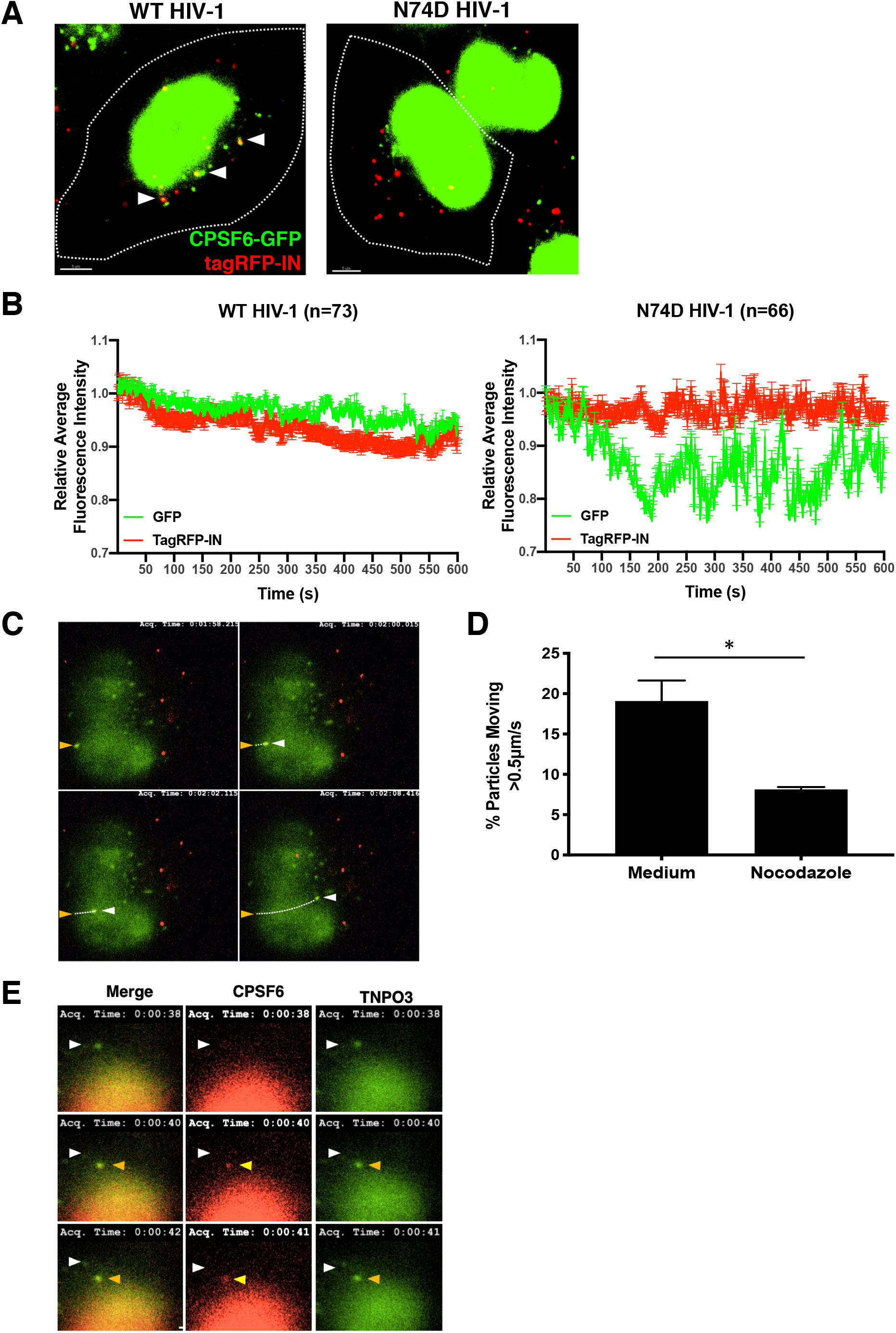
HIV-1 complexes associate with perinuclear CPSF6 and co-traffic on microtubules. (A) Confocal images of HeLa cells expressing CPSF6-GFP (green) are shown 60 min after infection with WT HIV-1 or N74D HIV-1 containing tagRFP-IN (red). White arrow indicate tagRFP-IN particles co-localized with CPSF6-GFP (yellow). (B) The average intensities of cytoplasmic tagRFP-IN from WT or N74D virus complexes and CPSF6-GFP during HILO imaging were measured for 10 min, normalized to 1.0 at time 0, and graphed. Error bars indicate STDEV. (C) The track of a WT HIV-1 mRuby3-IN complex (red) co-localized with CPSF6-GFP (green) is shown by HILO imaging in a HeLa cell over time (see Movie S3). The yellow arrow indicates the location of the complex at the first time point and white arrow indicates the location at subsequent time points. (D) The percentage of cytoplasmic CPSF6-GFP complexes co-localized with mRuby3-IN (n = 156-189) are shown that trafficked > 0.5 μm/s in HeLa cells treated with or without nocodazole. Error bars indicate standard errors of the mean (SEM). (E) A HeLa cell expressing GFP-TNPO3 (green) and CPSF6-iRFP (red) is shown. The white arrow indicates the location of a GFP-TNPO3 complex at the first time point that becomes co-localized with CPSF6-iRFP at subsequent time points (yellow arrow; see Movie S4).

### Changes in the CPSF6 RS domain alter WT HIV-1 complex trafficking

Truncation of CPSF6 at residue 358 (CPSF6-358), which removes the RS domain, or alteration of four positively charged amino acids (K547, R549, R559 and R561) in the RS domain to glutamic acid (CPSF6-4Glu) (**Fig 3A**) leads to decreased nuclear localization of CPSF6 and restriction of WT HIV-1 infection at the step of nuclear entry (Jang et al., 2019; Lee et al., 2010). Expression of iRFP670 tagged CPSF6-4Glu or CPSF6-358 led to increased cytoplasmic localization in cells, with the truncated protein showing a greater effect than the 4Glu mutant (**Fig 3B**). Similarly, expression of CPSF6-358 resulted in greater restriction of WT HIV-1 infection than expression of CPSF6-4Glu (**Fig 3C**). N74D HIV-1, which does not bind to CPSF6, was not restricted by either mutant.

**Figure 3.**
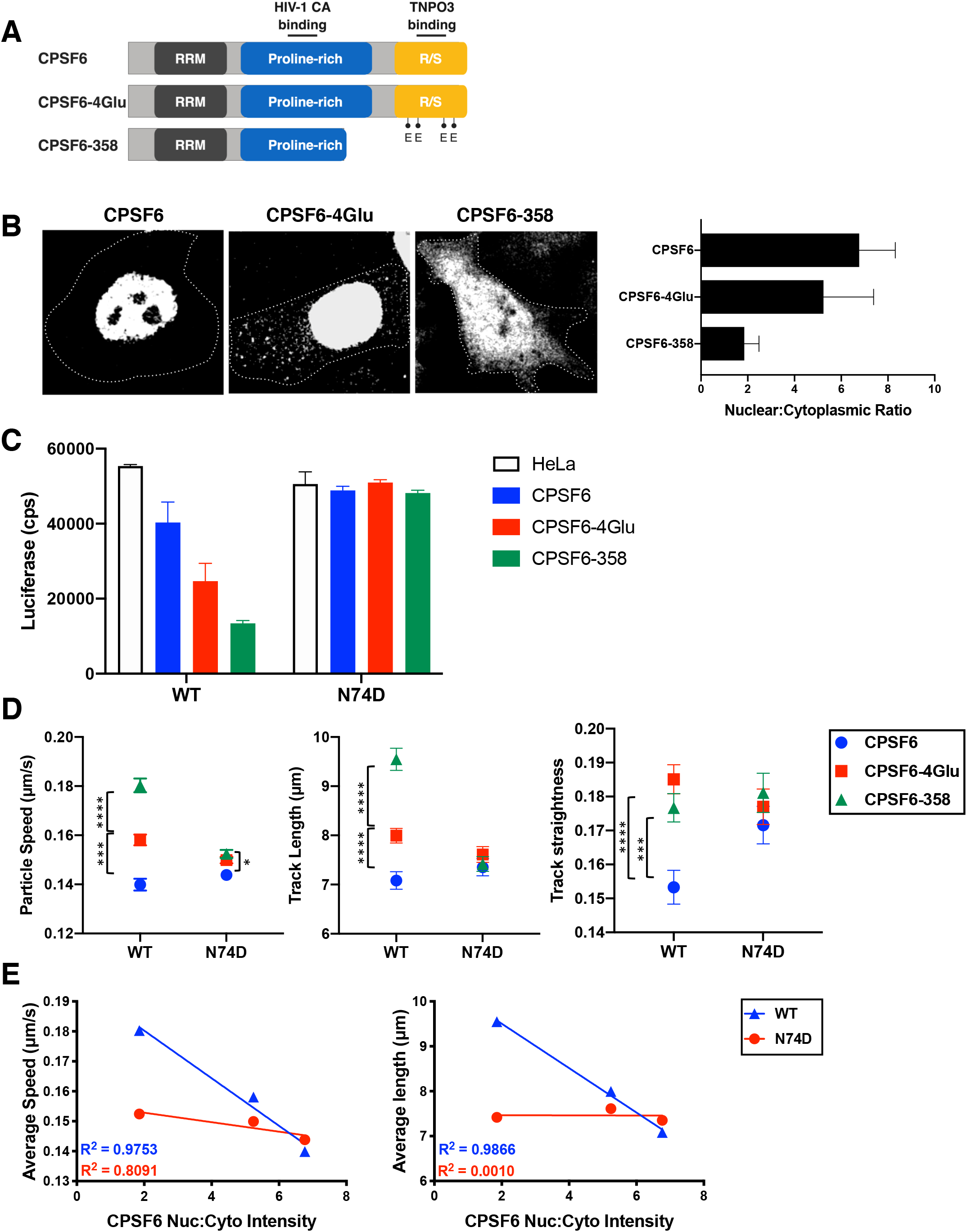
Alteration of the CPSF6 RS domain impacts HIV-1 trafficking. (A) CPSF6 protein domains and associated CPSF6-4Glu and CPSF6-358 changes. (B) Confocal microscopy images are shown of cells expressing CPSF6-iRFP, CPSF6-4Glu-iRFP, or CPSF6-358-iRFP. The cell periphery is indicated by dotted lines. The graph shows the nuclear to cytoplasmic ratio of CPSF6 for each. (C) Infection of WT and N74D HIV-1 in normal HeLa cells or HeLa cells expressing CPSF6-iRFP, CPSF6-4Glu-iRFP, or CPSF6-358-iRFP is shown as luciferase expression (counts per second, or cps) from a representative of 2 independent experiments. Error bars indicate STDEV of duplicates. (D) The average particle speeds, track lengths, and track straightness of WT or N74D HIV-1 mRuby3-IN complexes in HeLa cells expressing CPSF6-iRFP, CPSF6-4Glu-iRFP, or CPSF6-358-iRFP are shown from one of 2 independent HILO live-cell imaging experiments (see also Figure S2). Error bars indicate SEM. (E) Correlations of the average particle speed or average track length and CPSF6 nuclear:cytoplasmic ratio are shown for WT and N74D HIV-1.

To determine if relocalization of CPSF6 to the cytoplasm affects HIV-1 complex trafficking, live-cell imaging of WT and N74D HIV-1 complexes was performed in cells expressing fluorescently tagged CPSF6, CPSF6-4Glu, or CPSF6-358. To avoid imaging particles that have not yet fused out of endosomes into the cytoplasm, imaging was performed using mRuby3-IN labeled WT HIV-1 particles that were also labeled with the glycosylphosphatidylinositol targeting motif of decay-accelerating factor (Caras, 1991) tagged with fluorogen activating protein (FAP-GPI) to label the virus membrane. Loss of FAP-GPI signal from the mRuby3 signal signified that the HIV-1 membrane had fused with the endosome and the contents of the virus have been released into the cytoplasm (**Movie S5**). During synchronized infection, we observed that nearly all mRuby3 signal separated from the viral membrane by 50 min (**Fig S1**). Thus, for virus tracking experiments, acquisition of images began at 60 min post-infection.

In HeLa cells expressing CPSF6-GFP, WT and N74D viral complexes had similar speeds, and track lengths but differed in track straightness (**Figs 3D, S2**). In contrast, WT HIV-1 particles increased in all three measurements when CPSF6-4Glu-iRFP670 or CPSF6-358-iRFP670 was expressed in cells (**Figs 3D, S2**). Little or no change in particle speed, track length, or track straightness was observed for N74D complexes in the presence of CPSF6-4Glu or CPSF6-358. Similar to the effect of increased CPSF6 cytoplasmic localization on WT HIV-1 infectivity, average WT virus particle speed and track length inversely correlated with intensity of nuclear CPSF6 expression (**Fig 3E**). These data demonstrate that HIV-1 complex trafficking is altered in the cytoplasm via enhancing CPSF6 cytoplasmic localization. Furthermore, increased HIV-1 trafficking is associated with an infectivity defect.

### CPSF6 oligomerizes and disrupts assembled WT CA

Previously we showed that purified CPSF6-358 formed oligomers and disrupted WT HIV-1 CA tubular assemblies *in vitro* (Ning et al., 2018). To determine whether full-length CPSF6 had similar properties, it was purified and characterized with WT and N74D CA tubular assemblies. To obtain soluble CPSF6, an N-terminal maltose binding protein (MBP) fusion construct was expressed and purified, resulting in two peaks in size exclusion chromatography (**Fig S3A**, labeled P1 and P2) that corresponded to the tagged full-length CPSF6, as confirmed by western blot (**Figs S3B**). This suggests that the purified fusion protein may adopt different oligomeric states similar to what was observed for CPSF6-358, which also displayed two peaks in a size exclusion chromatography profile with dimer and large oligomers (Ning et al., 2018). Removal of the MBP-tag with HRV-3C protease resulted in precipitation of CPSF6 from both P1 and P2 (**Fig S3C**). Therefore, MBP-tagged soluble MBP-His_6_-CPSF6-588 (denoted as “MBP-CPSF6” henceforth) was used for further binding experiments.

Incubation of *in vitro* preassembled WT HIV-1 CA tubes with MBP-CPSF6 (both P1 and P2) resulted in co-sedimentation of MBP-CPSF6/CA complexes in the pelleted fractions (**Fig 4A**). Negligible binding of MBP-CPSF6 to N74D HIV-1 CA tubes was observed under the same assay conditions. TEM of the negative stained samples showed a drastic structural disruption of capsid tubes when incubated with MBP-CPSF6 (P1 or P2), while N74D CA tubes remained intact (**Fig 4C**). MBP-CPSF6, by itself, formed protein oligomers for both P1 and P2 fractions in CA assembly buffer (**Fig 4B**). Binding of MBP-CPSF6 to WT CA tubes resulted in dissolution of tubes and an appearance of distinct curved capsid remnants associated with MBP-CPSF6 densities. Intriguingly, the amount of pelletable capsid did not change upon capsid disruption (**Fig 4A**), suggesting that the predominant effect of MBP-CPSF6 is HIV-1 capsid fragmentation without dissociation into soluble proteins. Dose-dependent binding of MBP-CPSF6 to CA tubes was observed for both MBP-CPSF6 P1 and P2 by TEM (**Fig S4**). This represents the first direct evidence of full-length CPSF6 binding to and disruption of WT CA tube assemblies.

**Figure 4.**
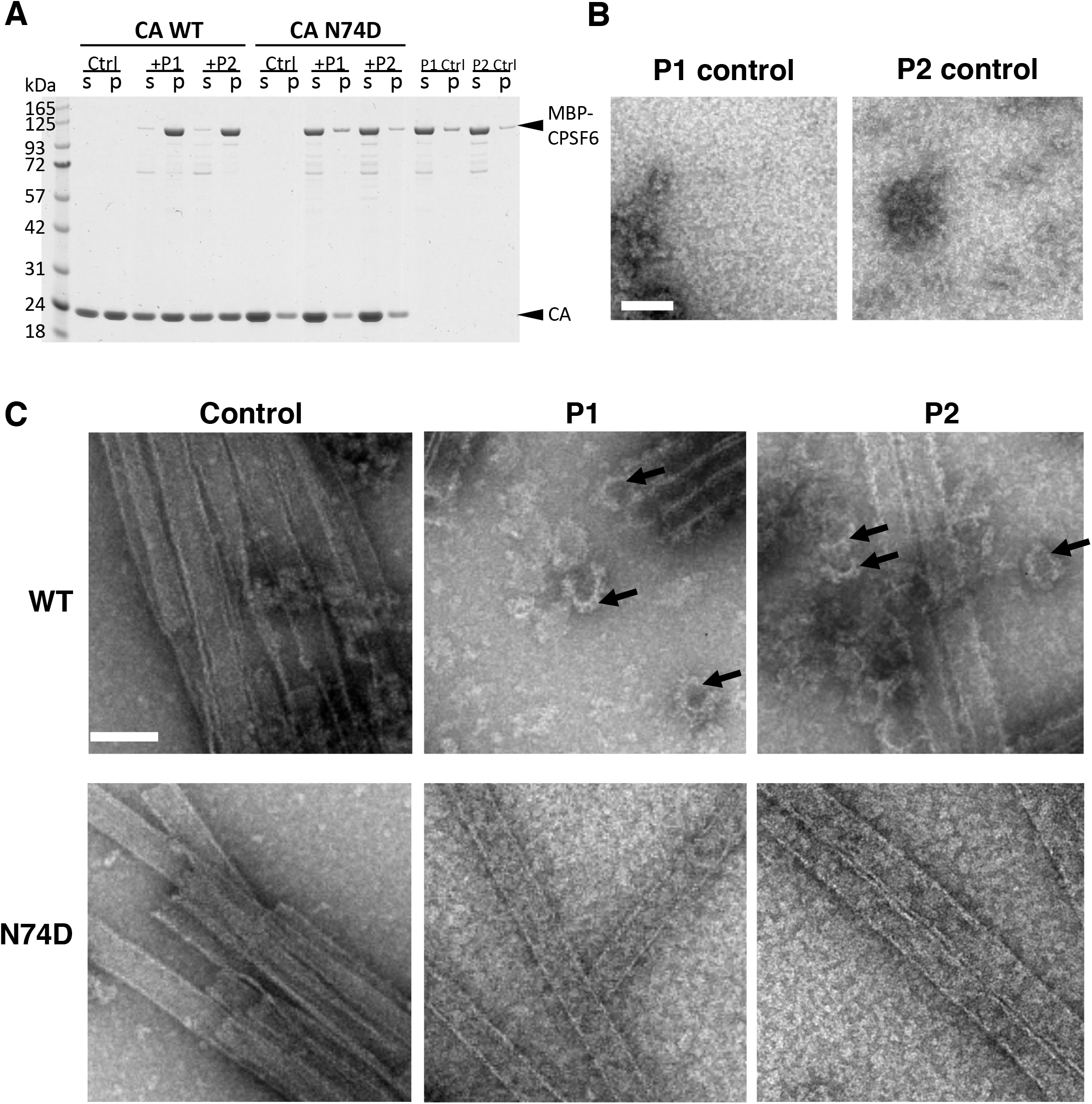
CPSF6 binds to and disrupts CA tubular assemblies. (A) SDS-PAGE analysis of WT and N74D CA assemblies following incubation with MBP-CPSF6 P1 or P2 and centrifugation. The gel was stained with Coomassie blue; supernatant (s) and pellet (p) samples are indicated. (B) Representative negative-stain EM micrographs are shown of P1 or P2 incubated in buffer. (C) Representative negative-stain EM micrographs are shown of WT CA (top panel; see also Figure S4) and CA N74D (bottom panel) tubular assemblies alone (control) or with 9 µM MBP-CPSF6 P1 or P2. The arrows indicate capsid fragments. Scale bars, 100 nm.

### HIV-1 induces cytoplasmic higher order CPSF6 complex formation in a CypA-dependent manner

In cells, CPSF6-358 forms puncta around WT HIV-1 mRuby3-IN complexes early after virus entry, leading to premature capsid permeabilization (Ning et al., 2018). These did not form in the presence of a small molecule inhibitor, PF74, which blocks CPSF6 binding to capsid (Matreyek et al., 2013), or if N74D HIV-1 was used. To determine if full-length higher order CPSF6 complexes form in the cytoplasm, immunostaining of CPSF6 was performed before and after HIV-1 infection. Higher order CPSF6 complexes were visualized in the perinuclear region after WT HIV-1 infection but not after N74D HIV-1 infection (**Figs 5A, 5B**), and were associated with IN-containing complexes (**Fig 2**) and CA (p24) staining (**Fig S5A**). CPSF6 puncta were also observed in the nuclei of cells at later time points after WT HIV-1 infection (**Fig S6**). Consistent with our *in vitro* results (**Fig 4**), these data suggest that CPSF6 binds to WT capsid in the cytoplasm of infected cells.

**Figure 5.**
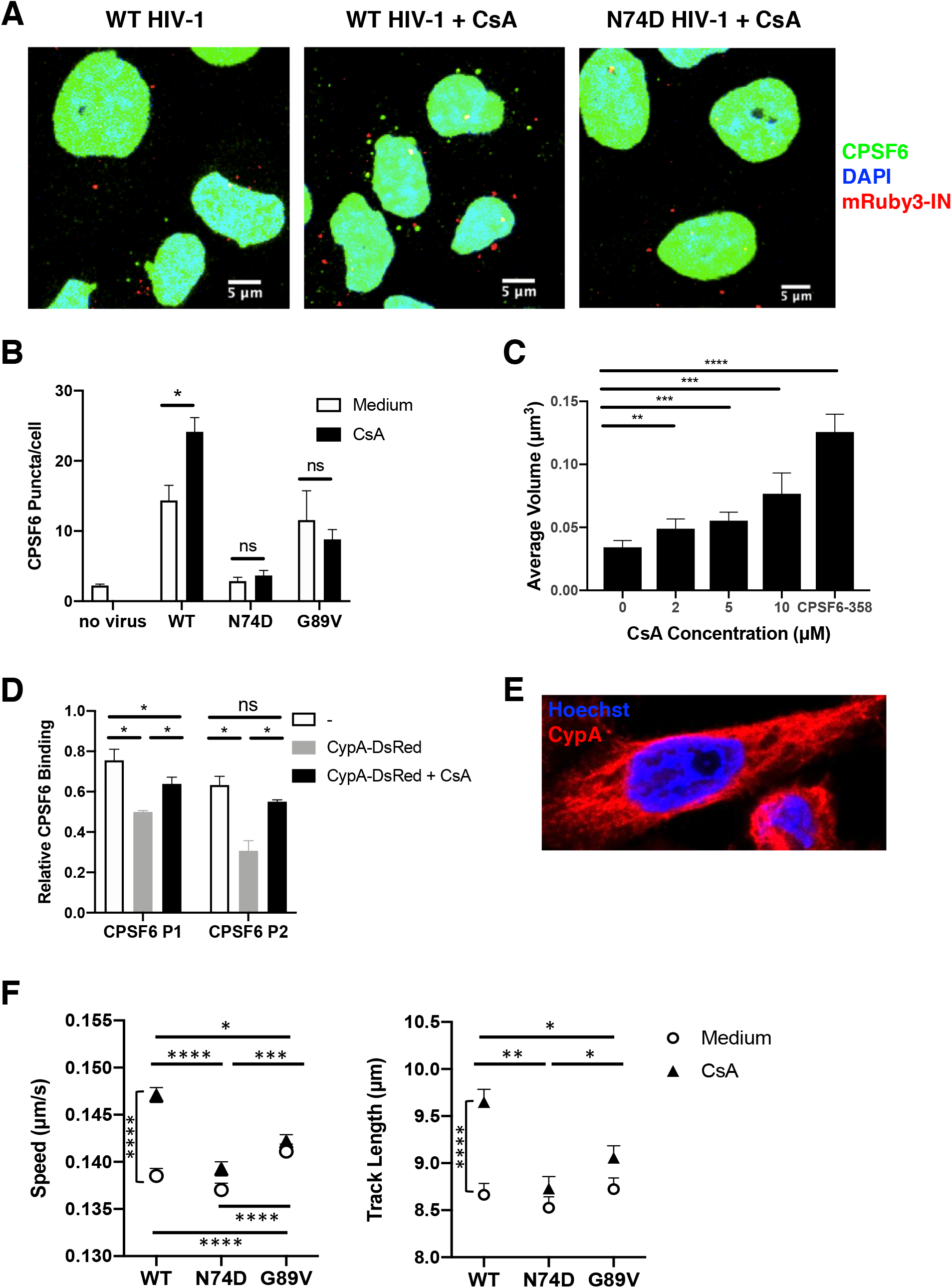
HIV-1 induction of cytoplasmic CPSF6 higher order complex formation as a function of CypA binding. (A) Representative confocal micrographs are shown of HeLa cells stained with CPSF6 antibodies (green) and Hoechst (blue) after infection with WT HIV-1 and WT or N74D HIV-1 in the presence of 10 μm CsA. Viruses contain mRuby3-IN (red). (B) Cytoplasmic CPSF6 staining was quantified in cells (n ≥ 1235) treated with or without 10 μm CsA and infected for 1 h with WT, N74D, or G89V HIV-1. (C) The average volume of cytoplasmic CPSF6 puncta (n ≥ 1235) was quantified in HeLa cells treated with or without CsA. Cells expressing CPSF6-358-GFP were used as a positive control. (D) CA-NC assemblies were incubated for 1 h in buffer, with CypA-DsRed, or CypA-DsRed and CsA, then incubated with MBP-CPSF6 P1 or P2 for 1 h. The relative binding of CPSF6 binding to CA-NC assemblies was quantified in each sample from stained SDS-PAGE gels. Error bars indicate STDEV from 3 independent experiments. (E) A representative confocal micrograph of a HeLa cell stained with Hoechst (blue) and CypA antibodies (red). (See also Figures S8 and S9). (F) The average particle speeds and track lengths of WT, N74D, or G89V HIV-1 mRuby3-IN complexes in HeLa cells treated with or without 10 μm CsA are shown for 1 of 3 independent experiments (see also Figure S10). Error bars indicate SEM.

Our previous work demonstrated that higher order CPSF6-358 complexes were larger and formed more rapidly when CypA binding to CA was inhibited by cyclosporine A (CsA) (Ning et al., 2018). Therefore, we examined whether the same would be true for full-length CPSF6. Indeed, when cells were treated with CsA and infected with WT HIV-1, greater numbers of CPSF6 higher order complexes were observed (**Figs 5A, 5B**). The volume of the complexes that formed in the presence of WT virus increased with increasing concentrations of CsA (**Fig 5C**), suggesting that inhibiting more CypA binding allowed more CPSF6 to bind to HIV-1 capsid. In contrast, CPSF6 complex formation was indistinguishable from background after infection with N74D HIV-1 (**Figs 5A, 5B**). To confirm that loss of CypA binding to capsid was responsible for the increase in CPSF6 higher order complexes, cells were infected with HIV-1 CA mutant G89V that is defective for CypA binding (Wiegers et al., 1999). Because G89V HIV-1 is restricted by CPSF6-358 and thus can still bind CPSF6 (De Iaco et al., 2013), we expected that this virus would induce CPSF6-GFP puncta that would not increase in number in the presence of CsA, which is what was observed (**Fig 5B**).

As removal of CypA from HIV-1 capsid enhanced CPSF6 higher order complex formation, we hypothesized that CPSF6 complex formation may be prevented if CypA binding to capsid was enhanced. Thus, virus was produced in the presence of CypA-DsRed, an oligomeric form of fluorescently labeled CypA with higher avidity to HIV-1 capsid than unlabeled CypA (Francis et al., 2016). Cells were infected with WT HIV-1 labeled with CypA-DsRed or with mRuby3-IN in the presence or absence of CsA and stained for CA (p24; **Fig S5B**). Similar levels of p24 staining were observed under all conditions. Virus containing mRuby3-IN led to formation of many CPSF6-358-GFP puncta associated with p24 that increased with CsA treatment (**Fig S5A**). However, cells infected with CypA-DsRed-labeled HIV-1 had significantly fewer GFP puncta and CsA treatment did not increase their formation, suggesting that enhanced CypA binding to capsid prevents HIV-1 interaction with CPSF6-358 and, likely, also CPSF6.

To directly test the ability of CypA to shield CPSF6 binding to HIV-1 capsid, MBP-CPSF6 protein (P1 and P2) binding to nanotubes composed of recombinant WT CA-SP1-nucleocapsid (CA-NC) protein was quantified in the presence or absence of CypA-DsRed, and in the presence or absence of cyclosporine A (CsA). (**Fig S7A**). Binding of CypA-DsRed to CA-NC tubes was not affected by subsequent MBP-CPSF6 binding but was inhibited by CsA treatment (**Fig S7B**). MBP-CPSF6 binding to HIV-1 CA-NC tubes decreased when CypA-DsRed was already bound, effects that were rescued partially (P1) or completely (P2) by the presence of CsA (**Fig 5D**). Collectively, these data demonstrate that CypA binding to capsid prevents CPSF6 from binding.

### Loss of CypA binding leads to altered cytoplasmic trafficking of WT HIV-1 complexes in a CPSF6-dependent manner

Although CypA is packaged into virions, HIV-1 capsid interactions with target cell CypA dictate infectivity (Sokolskaja et al., 2004). In contrast to CPSF6 expression, endogenous CypA localized to the cytoplasm of HeLa cells with somewhat of a filamentous appearance (**Figs 5E, S8**). Interestingly, not only was CypA excluded from the nucleus, its expression was absent from portions of the perinuclear region (**Fig S8**) that corresponded to the microtubule-organizing center (**Fig S9**).

To determine whether loss of CypA binding to HIV-1 capsid could affect virus trafficking, live-cell microscopy was performed on WT, N74D, and G89V HIV-1 in the presence and absence of CsA in HeLa cells (**Figs 5F, S10A**) or HeLa cells expressing CPSF6-GFP (**Figs S10B and S10C**). In the absence of drug, the average speed and track length of viral complexes were similar for WT HIV-1 and N74D HIV-1, while G89V HIV-1 complexes trafficked significantly faster and had similar track lengths. As observed for WT HIV-1 trafficking in cells expressing mutant CPSF6 (**Fig 5**), a higher rate of speed of G89V viral complexes was associated with lower infectivity (**Fig S11**). However, CsA treatment led to significantly increased speed and track length of WT HIV-1 particles. CsA did not affect N74D or G89V viral particles. These results suggest that loss of CypA binding to WT capsid influences trafficking of HIV-1 complexes in the cytoplasm in a CPSF6-dependent manner, as N74D viral particles were not affected by CsA treatment. G89V complexes that bind to CPSF6 but not CypA had altered trafficking and lower infectivity irrespective of CsA treatment. The expression and trafficking data together suggest that CypA prevents virus cores from binding prematurely to CPSF6 during trafficking to the nucleus.

### Depletion of CPSF6 rescues HIV-1 complex trafficking defect caused by loss of CypA binding

To validate that the effect of CsA treatment on WT viral complex trafficking is mediated by CPSF6, CPSF6 was depleted from cells using shRNA knockdown. HeLa cells were transduced with lentiviruses expressing a shRNA targeting CPSF6 or a scrambled control shRNA. Knockdown (KD) of CPSF6 was verified by immunofluorescence staining (**Fig 6A**). In the absence of CsA, CPSF6 KD had no effect on WT virus particle tracking (**Fig 6B**). Infectivity of WT HIV-1 in untreated cells decreased after CPSF6 depletion (**Fig 6C**), which may be attributable to previously described reduced cell proliferation of CPSF6 knockout cells (Sowd et al., 2016). As shown above (**Fig 5F**), CsA led to a significant increase in WT HIV-1 complex speed and track length in HeLa cells (**Figs 6B, S12**). However, CPSF6 KD led to decreased WT HIV-1 particle speed to that of unmodified, untreated HeLa cells.

**Figure 6.**
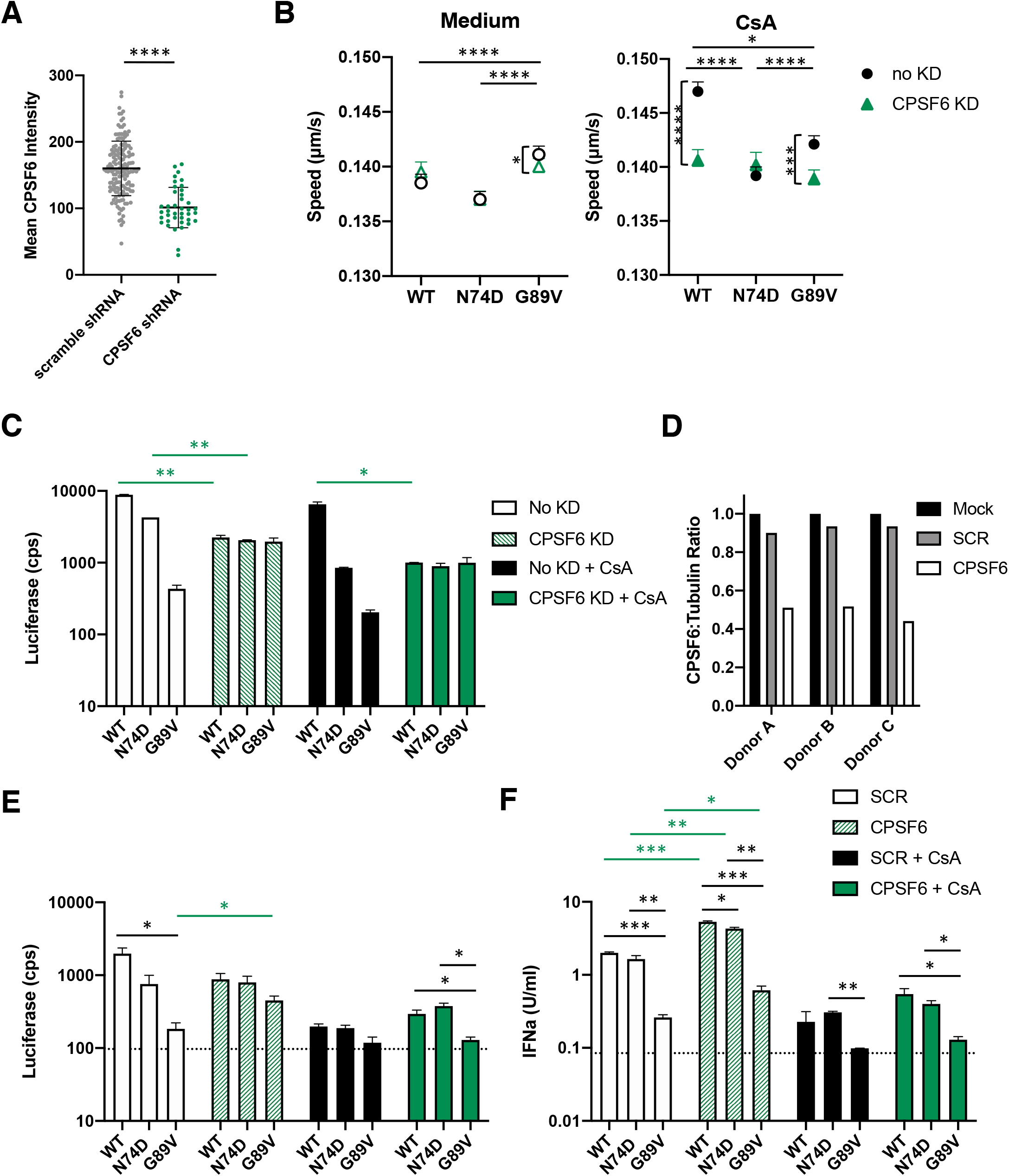
CPSF6 knockdown in HeLa cells and primary PBMC alters HIV-1 trafficking and decreases infection in the absence of CypA binding. (A) The maximum intensity of CPSF6 antibody staining is shown from individual HeLa cells transduced with lentiviruses expressing scrambled (SCR) or CPSF6 shRNA. The error bars indicate STDEV (n ≥ 157 cells). (B) The average particle speeds of WT, N74D, or G89V HIV-1 mRuby3-IN complexes in HeLa cells expressing SCR or CPSF6 shRNA and treated with or without 10 μm CsA are shown. Representative of 2 independent experiments (see also Figure S12). Error bars indicate SEM. (C) The infection of WT, N74D, or G89V HIV-1 in the same cells shown in (B) are graphed. Error bars indicate STDEV of duplicates. (D) CPSF6 depletion was quantified by the ratio of CPSF6 to α-tubulin measured by western blot for activated PBMCs from each donor. (See also Figure S13A.) (E) Representative infections with WT, N74D, and G89V HIV-1 were measured in primary PBMC cells from a donor after shRNA depletion of CPSF6, treatment with CsA, or both. (See also Figure S13B.) (F) IFNα production from infected PBMC shown in (E) was quantified using HEK 293T ISRE-luc indicator cells. (See also Figure S13C.)

The CA mutants showed different cytoplasmic trafficking patterns compared to WT HIV-1, while N74D complex trafficking was unaffected by CPSF6 KD and CsA treatment (**Fig 6B**). Interestingly, depletion of CPSF6 led to a significant decrease in G89V HIV-1 complex trafficking with or without CsA treatment (**Fig 6B**), which corresponded to a rescue of the infectivity defect of this mutant (**Fig 6C**). Our data indicate that CypA alters HIV-1 trafficking in a CPSF6-dependent manner, further suggesting that CypA binding protects HIV-1 capsid from binding too prematurely to or too much of CPSF6.

### Depletion of CPSF6 rescues G89V HIV-1 infectivity and induces interferon (IFN) α production in primary PBMC

Infectivity of WT HIV-1 and CA mutants were performed in phytohemagglutinin (PHA)-stimulated primary human peripheral blood mononuclear cells (PBMC) from 3 donors. Cells were infected in the presence or absence of CPSF6 depletion and CsA treatment. Depletion of CPSF6 was verified by western blot quantification (**Figs 6D, S13A**). Similar to the HeLa cell data, infection of N74D and G89V HIV-1 was inhibited in primary PBMC (**Figs 6E, S13B**), as previously reported (Lee et al., 2010; Sokolskaja et al., 2006). Depletion of CPSF6 partially rescued G89V HIV-1 infectivity (**Fig 6E, S13B**), similar to what was observed in HeLa cells (**Fig 6C**).

It has been suggested that a loss of HIV-1 capsid interaction with host factors, such as CPSF6 and CypA, can lead to induction of type I interferon in immune cells (Rasaiyaah et al., 2013). Thus, IFNα production was quantified after HIV-1 infection from the supernatants of PBMC depleted of CPSF6 and/or treated with CsA. WT and N74D HIV-1 induced similar levels of IFNα after infection of CD4+ T cells, whereas G89V HIV-1 did not (**Fig 6F, S13C**). Depletion of CPSF6 significantly increased IFNα production after infection by all three viruses, consistent with previous results (Rasaiyaah et al., 2013). Treatment with CsA further inhibited IFNα production after infection with the viruses. Overall, IFNα production in primary CD4+ T cells correlated with the infectivity data, such that higher infection levels led to greater IFNα production.

### Depletion of CPSF6 in macrophages results in decreased HIV-1 infectivity independent of CypA binding

As we and others previously showed that N74D HIV-1 has an early infectivity defect in monocyte-derived macrophages (MDM) (Ambrose et al., 2012; Schaller et al., 2011), we investigated whether intracellular trafficking of HIV-1 is dependent on CPSF6 and CypA. MDM were stained for endogenous CypA and CPSF6 expression. As in HeLa cells, CPSF6 expression was nuclear in MDM (**Fig S14**). However, infection with WT HIV-1 failed to induce higher order CPSF6 complex formation in the cytoplasm (data not shown), consistent with previous results (Bejarano et al., 2019) and indicative of less cytoplasmic CPSF6 expression in MDM compared to HeLa cells. In contrast, CypA expression differed greatly between cell types. MDM had pronounced nuclear CypA expression in addition to patches of plasma membrane and cytoplasmic expression (**Figs 7A, S15**).

**Figure 7.**
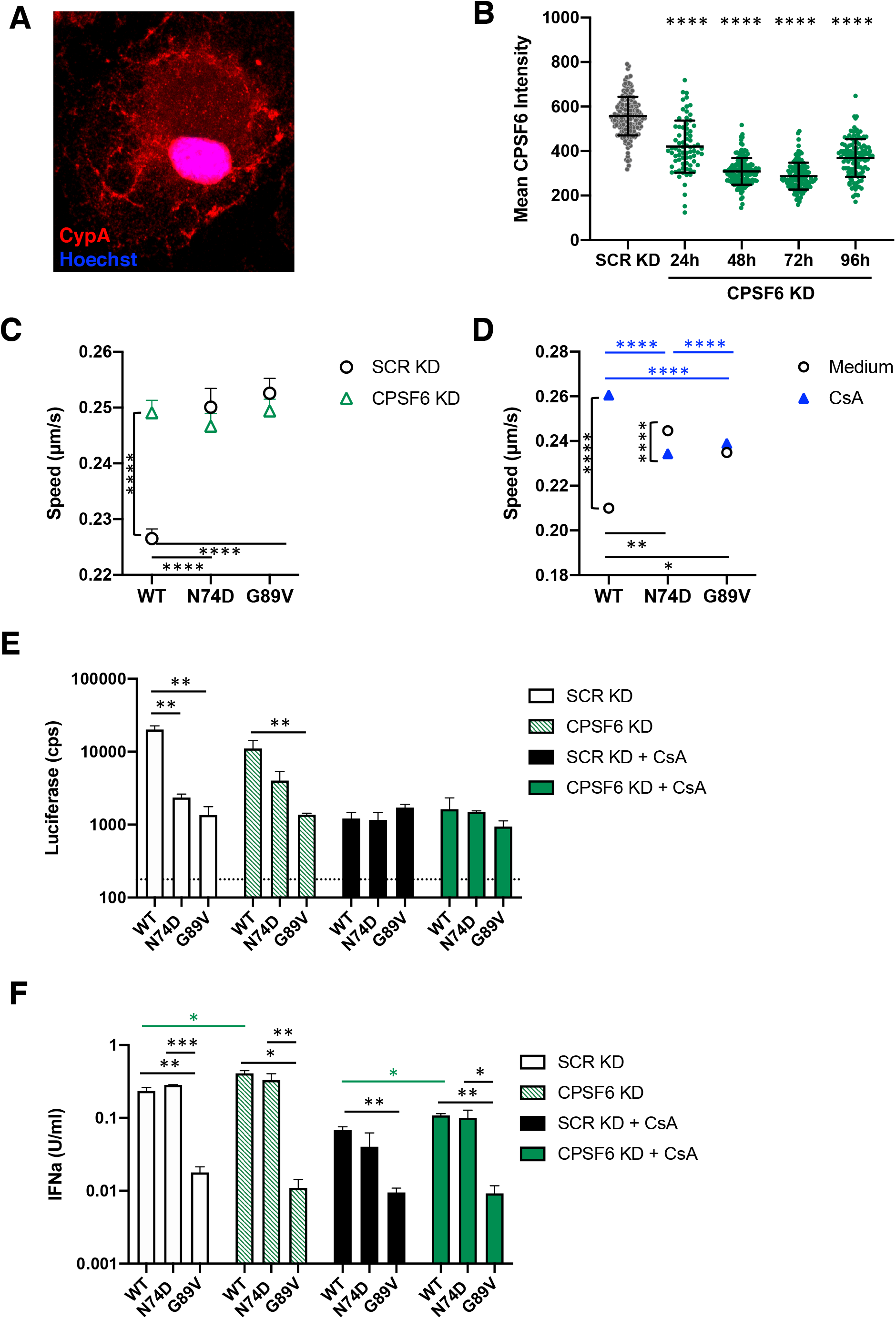
CPSF6 KD decreases HIV-1 infection in MDM and enhances IFNα. (A) A representative confocal micrograph of a MDM stained with Hoechst (blue) and CypA (red) is shown. (Also see Figure S15.) (B) The mean intensity of CPSF6 antibody staining is shown from individual MDM transduced with lentiviruses expressing SCR or CPSF6 shRNA. The error bars indicate STDEV of n ≥ 77 cells. (See also Figure S14.) (C) The average particle speeds of WT, N74D, or G89V HIV-1 mRuby3-IN complexes in MDM from 1 of 3 individual donors with SCR KD or CPSF6 KD are shown (see also Figure S16). Error bars indicate SEM. (D) The average particle speeds of WT, N74D, or G89V HIV-1 mRuby3-IN complexes in MDM from a representative donor treated with or without CsA are shown. Error bars indicate SEM. (E) The infection of WT, N74D, or G89V HIV-1 in the same cells shown in (B) are graphed (see also Figure S17A). Error bars indicate STDEV of duplicates and the dotted line represents the average luciferase expression of uninfected cells. (F) IFNα was measured in the supernatants of cells shown in (E; see also Figure S17B). Error bars indicate STDEV of duplicates. The limit of detection is 0.001 U/ml.

CPSF6 was depleted with shRNA in MDM, which was confirmed by immunostaining (**Figs 7B, S14**). Trafficking of mRuby3-IN complexes was evaluated in MDM from 3 donors with and without CPSF6 depletion. CPSF6 depletion led to a significant increase speed and track length of WT HIV-1 complexes (**Figs 7C, S16**). Loss of CPSF6 led to a modest decrease in WT HIV-1 single-cycle infectivity with only Donor 2 being significant (**Figs 7E, S17A**). However, CPSF6 depletion led to a significant decrease in spreading infection with WT HIV-1 but not with N74D HIV-1 in MDM (**Fig S17B**). Despite effects on infectivity, CPSF6 KD resulted in significantly increased IFNα production in all 3 donors (**Figs 7F, S17C**). In normal MDM, both N74D and G89V HIV-1 had significantly decreased infectivity which corresponded to faster mRuby3-IN trafficking. CPSF6 depletion had no effect on the trafficking of mutant CA complexes or subsequent infection. Interestingly, despite having low infectivity in MDM, N74D HIV-1 led to high IFNα production in MDM with or without CPSF6 KD, similar to WT HIV-1 infection (**Figs 7F, S17C**). Increased type I interferon production by N74D HIV-1 during reverse transcription in macrophages has been described previously (Rasaiyaah et al., 2013). G89V HIV-1 resulted in low infection and IFN production. These results suggest that loss of CPSF6 in macrophages affects WT HIV-1 CA trafficking and increased production of type I IFN, leading to decreased infectivity.

CsA treatment of MDM significantly increased WT HIV-1 complex trafficking speed (**Fig 7D**) with a corresponding decrease in infection in all donors (**Figs 7E, S17A**). The low infectivity of WT virus corresponded to a decrease in IFNα (**Figs 7F, S17C**). CPSF6 KD with CsA treatment (i.e. loss of CPSF6 and CypA binding) led to a significant increase in infectivity of WT virus in all donors with an increase of IFNα production compared to CsA treatment alone. N74D HIV-1 infection was significantly reduced during CsA treatment, with or without CPSF6 KD, but still resulted in higher IFNα. Although loss of CypA binding significantly reduces virus infectivity, these results suggest that alteration of HIV-1 trafficking, infectivity, and type I IFN response by depletion of CPSF6 is independent of CypA binding in macrophages.

## Discussion

HIV-1 capsid has been shown to interact directly or indirectly with many host cell factors (Yamashita and Engelman, 2017). Included in this long list is CPSF6, which is involved in mRNA splicing and adenylation in the nucleus (Lee et al., 2010). In this study, we detected cytoplasmic, punctate expression of endogenous CPSF6 and fluorescently tagged CPSF6 in the perinuclear region of cells. This is consistent with a recent study in which CPSF6 was associated with HIV-1 complexes in the cytoplasm prior to nuclear import (Burdick et al., 2020). Full-length MBP-CPSF6 protein forms oligomers *in vitro* that are capable of binding to HIV-1 CA assemblies, similar to what we previously reported for CPSF6-358 (Ning et al., 2018), which lacks the RS domain but retains the central proline-rich domain that mediates CA binding. In cells, CPSF6 puncta likely also represent higher-order complexes that bind to HIV-1 capsid in the cytoplasm after viral entry. Removal or truncation of the CPSF6 RS domain leads to mislocalization of CPSF6 to the periphery of the cell due to loss of TNPO3 binding, and reduced HIV-1 nuclear import and infectivity (De Iaco et al., 2013; Jang et al., 2019; Lee et al., 2010). Here we demonstrate with live-cell imaging that HIV-1 complexes traffic with CPSF6 in a capsid-dependent manner. In addition, virus complexes increase in speed in the presence of CPSF6 RS mutants or by introducing the CA N74D mutation that abolishes CPSF6 binding. Interestingly, increased microtubule trafficking of HIV-1 is associated with reduced infectivity. Previously, we showed significant colocalization of CPSF6-358 to mRuby3-IN complexes in the cytoplasm, likely due to high concentrations at the cell periphery that is not seen with full-length CPSF6 (Ning et al., 2018). It is possible that faster or more efficient trafficking is due to more capsid-bound CPSF6, which may alter accessibility of capsid to certain host proteins, such as microtubule motor proteins or motor adaptors.

Previously it was shown that HIV-1 capsid traffics on microtubules on its way to the nucleus (Dharan and Campbell, 2018). HIV-1 capsid uncoating is delayed by destabilization of microtubules or knockdown of microtubule motor proteins kinesin and dynein or NPC protein Nup358 (Dharan et al., 2016; Lukic et al., 2014; Malikov and Naghavi, 2017), suggesting that microtubule trafficking and NPC binding is linked to capsid uncoating. Recent work from several groups suggests that the main uncoating event occurs at or inside the nucleus (Arhel et al., 2007; Burdick et al., 2017; Burdick et al., 2020; Francis and Melikyan, 2018; Zurnic Bonisch et al., 2020). Here we demonstrate that HIV-1 capsid traffics with CPSF6 and TNPO3 on microtubules and that CPSF6 facilitates CA tubular disassembly. Our previous work demonstrated that CPSF6-358 associates with HIV-1 complexes in a CA-dependent manner and leads to more rapid uncoating kinetics and reduced virus infectivity (Ning et al., 2018). Therefore, CPSF6 may play a role in both HIV-1 capsid uncoating, which may initiate in the cytoplasm, and nuclear import (Bejarano et al., 2019; Chin et al., 2015).

CypA binding to HIV-1 capsid was described nearly three decades ago (Luban et al., 1993). Yet until recently its role in HIV-1 infection was ill defined. Here we show that loss of CypA binding to HIV-1 capsid in infected cells due to CsA treatment leads to increased capsid binding to CPSF6, which is consistent with our previous results with fluorescently labeled CPSF6-358 (Ning et al., 2018). Conversely, production of HIV-1 in the presence of CypA-DsRed, which has increased binding to capsid compared to untagged CypA (Francis et al., 2016), reduces CPSF6 binding to capsid. This is also seen *in vitro* in a competitive binding assay with CA-NC assemblies in the presence of CypA-DsRed and MBP-CPSF6 proteins. Trafficking of HIV-1 complexes in cells increases in the presence of CsA or with the G89V CA mutation, which is again correlated with decreased infectivity. These results suggest that CypA binding to capsid prevents CPSF6 binding. As CypA expression in HeLa cells is seemingly restricted to the cytoplasm, which is where CPSF6 expression is excluded, we hypothesize that CypA interacts with HIV-1 capsid in the cell periphery first and prevents CPSF6 binding to HIV-1 capsid before it is near the nucleus, ensuring that uncoating does not occur prematurely.

Depletion of CPSF6 in cells does not affect HIV-1 infection in HeLa cells or PBMC, as previously demonstrated (Lee et al., 2010), nor does it alter HIV-1 trafficking to the nucleus in HeLa cells. As N74D HIV-1 is associated with microtubules and traffics in a linear fashion like WT HIV-1, it suggests that HIV-1 microtubule trafficking and nuclear import can be independent of CPSF6. Surprisingly, loss of CPSF6 expression does affect WT and G89V HIV-1 trafficking during CsA treatment but not N74D HIV-1 trafficking. Alterations in HeLa trafficking under these conditions correspond with HeLa infectivity data. These results again highlight the interplay of CPSF6 and CypA in capsid interaction. As it has not been possible to knockout CPSF6 or completely deplete CPSF6 in these cells, these data further suggest that loss of CypA binding allows CPSF6 binding to occur outside of the nucleus to affect trafficking and infectivity.

Although high-speed imaging of HIV-1 particles in CD4+ T cells was not possible, infection results after CPSF6 depletion and/or CsA treatment in primary PBMC largely mimic those seen in HeLa cells. However, CsA treatment significantly reduces infectivity of WT HIV-1 as well as the CA mutants. These results correlate with IFNα production. A recent report shows that loss of CypA binding, either by depletion, CsA treatment or CA mutations, leads to significant restriction by human TRIM5α in CD4+ T cells (Selyutina et al., 2020). Thus, CypA binding to HIV-1 capsid may be protective against multiple capsid-binding cellular factors.

Capsid has been shown to play an important part in infection of non-dividing cells such as macrophages (Yamashita and Emerman, 2009). CPSF6-independent HIV-1 (e.g., N74D CA mutant) and mutants that do not bind to CypA (e.g., P90A CA mutant) are restricted in MDM prior to or at initiation of reverse transcription (Ambrose et al., 2012; Kim et al., 2019). Here we show that trafficking of N74D and G89V HIV-1 complexes in the cytoplasm is aberrant in MDM. Depletion of CPSF6 or treatment with CsA in MDM also lead to improper trafficking of WT HIV-1 complexes, which corresponds with reduced infectivity. Interestingly, WT HIV-1 infection of MDM with CPSF6 knockdown or N74D HIV-1 infection of MDM leads to increased IFNα production, consistent with a previous study showing that CPSF6 plays a role in immune sensing in MDM (Rasaiyaah et al., 2013). As in CD4+ T cells, loss of CypA binding to HIV-1 capsid leads to human TRIM5α restriction prior to completion of reverse transcription (Kim et al., 2019).

Differences in capsid uncoating and PIC composition have also been observed in MDM compared to other cell types. For example, cytoplasmic HIV-1 DNA staining was nearly 100% colocalized with CA staining in TZM-bl cells whereas there was less than 50% colocalization in MDMs (Peng et al., 2014). Whereas most CA staining is lost from PICs in the nucleus of HeLa cells, most nuclear PICs in MDMs were CA positive (Chin et al., 2015; Peng et al., 2014) and colocalized with CPSF6 puncta (Bejarano et al., 2019). Here we show that localization of CypA in MDM is both cytoplasmic and nuclear, whereas it is only expressed in the periphery of the cell in HeLa cells. Also, little to no cytoplasmic CPSF6 puncta are detected in MDM. As we show that CPSF6 binding to capsid is prevented by CypA binding and promotes capsid dissociation, CypA expression in the perinuclear region and nucleus of MDM could explain why more CA remains associated with the viral genome in these nuclei as compared to HeLa cells. Our results contribute to the growing literature of the ability of HIV-1 capsid to bind multiple host cell factors in a highly orchestrated manner to promote viral infectivity, which differs depending on the specific cellular environment.

## Materials and Methods

### Plasmids

The replication-defective HIV-1 proviral plasmid pNLdE-luc (WT and mutants) has been described (Ambrose et al., 2012; Lee et al., 2010). Viruses were pseudotyped with pL-VSV-G (Lee and Linial, 1994). N74D was also introduced similarly in a replication-competent proviral plasmid pNL4-BAL (gift from Ned Landau). pVpr-mRuby3-IN was previously described (Ning et al., 2018) and a similar version, pVpr-tagRFP-IN was also used.

Lentiviral vectors were used to introduce tagged host proteins or shRNA to cells. Fluorescently tagged CPSF6 and related mutants were introduced into the pSICO vector using BamHI and NotI sites. pSICO-GFP-TNPO3 was created using same sites with GFP-TNPO3 (gift from Ned Landau). pLenti-FAP-GPI was created using the BamHI and NotI sites to move the FAP-GPI sequence from pcDNA-IgKappa-myc-dL5-2XG4S-GPI (Addgene 153308) into pLenti-puro (gift from Ie-Ming Shih; Addgene #39481). A plasmid encoding CypA-DsRed (Francis et al., 2016) was a gift from Greg Melikyan. The lentiviral vector pHIVSIREN expressing CPSF6 shRNA (Rasaiyaah et al., 2013) was a gift from Greg Towers. pLKO.1-puro-shNT (gift from Jacob Corn, Addgene #109012) was used as a scrambled shRNA control. The lentiviral packaging plasmid psPAX2 (gift from Dr. Didier Trono, Addgene #109012) and pCMV-VSV-G were used to produce viruses.

The *CPSF6* gene and the MBP tag were amplified by PCR and subcloned into the pcDNA3.1(+) mammalian expression vector (Thermo Fisher Scientific) using the NEBuilder HiFi Assembly kit (New England Biolabs) after linearization with the restriction enzymes EcoRV and Xba I. The resulting insert, designated as MBP-His_6_-CPSF6-588, has a leading Kozak sequence, an N-terminal MBP tag, followed by a hexahistidine tag (His_6_).

HIV-1 CA and CA-NC were previously described (Byeon et al., 2009; Meng et al., 2012). In brief, they were cloned from the cDNA of *Pr55^Gag^*, which was obtained from the NIH AIDS Research and Reference Reagent Program, Division of AIDS, NIAID, NIH. Briefly, CA and CA-NC regions were amplified and subcloned into pET21 (EMD Chemicals Inc.) using NdeI and XhoI sites. Proteins were expressed and purified as previously described for Gag (∆MA_15–100_∆p6) (Ning et al., 2016; Zhao et al., 2013). CypA-DsRed-Exp2 (gift from Greg Melikyan) was cloned into the pET28 vector, resulting in a N-terminal His_6_-tagged protein.

### Cells

HeLa and HEK 293T cell lines were cultured in Dulbecco’s modified Eagle medium (DMEM; Thermo Fisher Scientific) supplemented with 10% fetal bovine serum (FBS; Atlanta Biologicals), 100 U/ml penicillin, 100◻μg/ml streptomycin, and 2◻mM L-glutamine (PSG; Thermo Fisher Scientific) at 37° C, 5% CO_2_. HEK 293T cells stably expressing the firefly luciferase gene downstream of the interferon-sensitive response element (HEK293-ISRE-luc) (Larocque et al., 2011) were a gift from Rahm Gummuluru and were cultured in complete DMEM supplemented with 2◻μg/ml puromycin (Invitrogen). Stable cell lines were made by transduction with lentiviruses expressing fluorescently tagged host proteins followed by fluorescence activated cell sorting. GHOST-R3/X4/R5 lentiviral reporter cells (Cecilia et al., 1998) were cultured in DMEM supplemented 10% FBS, PSG, 100◻μg/ml geneticin G418 (Thermo Fisher Scientific), 0.5◻μg/ml puromycin (Invitrogen), and 100◻μg/ml hygromycin B (Invitrogen).

Human PBMCs were isolated from leukapheresis obtained from the Central Blood Bank (Pittsburgh, PA) using Ficoll-Paque Plus (GE Healthcare) density gradient centrifugation, following the manufacturer’s instructions. PBMCs were cultured in RPMI 1640 medium (Thermo Fisher Scientific) supplemented with 10% FBS, PSG, and 20 U/ml recombinant interleukin-2 (IL-2; Thermo Fisher Scientific) at 37° C, 5% CO_2_. To expand T lymphocytes, PBMCs were stimulated with 50 U/ml IL-2 and 5◻μg/ml phytohemagglutinin (PHA; Sigma-Aldrich) for 72 h prior to infection or transduction. CD14+ monocytes were isolated from PBMCs using human anti-CD14 magnetic beads with LS Columns (Miltenyi Biotec). CD14+ monocytes were differentiated into MDM in RPMI 1640 medium supplemented with 10% FBS, PSG, and 50 ng/ml recombinant granulocyte–macrophage colony-stimulating factor (R&D Systems) for 7 days at 37° C, 5% CO_2_ prior to experimentation.

### Viruses

Replication-defective HIV-1_NL4-3_-luciferase virus pseudotyped with VSV-G was produced by transfection of HEK 293T cells or HEK 293T cells stably expressing FAP-GPI with pNLdE-luc, pL-VSV-G, and pVpr-pcs-mRuby3-IN/pVpr-tagRFP-IN/CypA-DsRed at a weight ratio of 5:5:1. Replication-competent HIV-1 was produced by transfection of HEK 293T cells with the full-length proviral construct. Lentiviruses encoding fluorescent host proteins were produced by transfecting HEK 293T cells with lentiviral, packaging, and pCMV-VSV-G plasmids at a weight ratio of 4:3:1. Transfections were performed using Lipofectamine 2000 (Invitrogen). Viruses were filtered through a 0.45 μM filters, concentrated with Lenti-X (Takara Bio) following the manufacturer’s protocol, and stored at −80° C. Viruses were quantified by p24 ELISA (XpressBio) and titered on GHOST-R3/X4/R5 cells. FAP-GPI, mRuby3-IN, and CypA-DsRed labeled viruses were assessed for labeling efficiency by total internal reflection fluorescence (TIRF) imaging.

### HIV-1 infection assays

HeLa cells and differentiated macrophages were seeded in 24-well plates overnight and then transduced with shRNA encoding viruses. 48 h post transduction, cells were infected with equal p24 amounts of luciferase reporter viruses for 48 h. Cells were lysed and assessed for luciferase production (Promega) with a 1450 MicroBeta TriLux microplate luminescence counter (PerkinElmer). CPSF6 KD efficiency was assessed by CPSF6 antibody staining (NBP1-85676). PHA-stimulated PBMCs were transduced with viruses encoding shRNA and selected with 2 μg/mL puromycin for 72 h. KD efficiency was measured by western blot. PBMCs were re-stimulated with PHA and challenged with luciferase reporter viruses for 72 h prior to luciferase measurement. For assays including treatment with CsA, cells were treated with CsA (10◻μM) at the time of plating and remained in drug-containing medium throughout the assay.

Replication of HIV-1 in macrophages was performed in duplicate by infecting transduced macrophages with WT or N74D HIV-1_NL4-BAL_ at a multiplicity of infection of 0.1. Supernatant was collected and new medium was added every two days. Viral replication was quantified by p24 production in the supernatant by ELISA (XpressBio) at day.

### Fluorescence microscopy

For fixed cell imaging, HeLa cells or macrophages were plated in MatTek dishes overnight. Synchronized infections were performed by incubation at 4° C for 10 min, followed by aspiration of medium, addition of cold fluorescently labeled HIV-1 (5 ng p24), and further incubation at 4° C for 15 min to allow virus attachment. Cells were then incubated at 37° C for 20 min, followed by washing with warm medium and incubation in fresh medium. At 1 h post-infection, cells were washed with phosphate buffered saline, pH 7.4 (PBS) and fixed with 2% paraformaldehyde (PFA). After permeabilization with 0.1% TritonX-100 for 15 min, the fixed sample was blocked with serum matching the secondary antibody for 45 min. Primary antibodies were added to the fixed cells in PBB buffer (2% bovine serum albumin in PBS) for 1 h and washed with PBB. Secondary antibodies were added to the cells in PBB buffer for 1 h. After washing with PBB and PBS, the cells were stained with Hoechst (1:2000) and mounted with a coverslip using gelvetol.

A Nikon Ti inverted confocal microscope was used to acquire 3D stacks images of fixed samples with a 100X 1.49 NA oil-immersion objective. LU-NV laser launch (Nikon) was used to emit lasers at 405 nm, 488 nm, 561 nm, and 640 nm. Fields of view were randomly chosen by quick scanning in the Hoechst channel. ND Acquisition in Elements (Nikon) was applied to collect 3D multi-channel imaging (1024 x 1024 pixels) with 2X line averaging. Images of 488 nm and 561 nm channels were acquired by GaAsP detectors (Nikon). 3D stacks were acquired with 0.5 μm step intervals to cover the entire cell volume (6-10 μm) with a motorized piezo Z stage (Nikon).

For live-cell HILO imaging, a Nikon Ti TIRF microscope with a 100X 1.49 NA oil-immersion objective and a Photometrics Prime 95B sCMOS camera was used. In multi-color live-cell imaging experiments, a FLI high speed filter wheel was used. Synchronized infections in HeLa cells or macrophages were performed as described above. After shifting to 37° C for 20 min, cells were washed with pre-warmed fresh medium: FluoroBrite medium (Thermo Fisher Scientific) for HeLa cells or RPMI 1640 medium for macrophages. After 1 h post-infection, the MatTek dish was loaded on the stage insert and maintained at 37° C (Tokai Hit stage chamber). Each image was acquired at least 1 frame per second (FPS) to track viruses for 10 min. For visualizing microtubules, 1 μM SiR-tubulin (Cytoskeleton) was added to the medium 30 min prior to imaging. For visualizing viral membranes, the MG-B-Tau FAP dye (Perkins and Bruchez, 2020) was added to the virus at 500 nM for 10 min prior to addition to cells. RAM capture in Nikon Elements was used to achieve faster multi-color live-cell imaging (≥ 2FPS).

### Imaging quantification and data analysis

All imaging quantification was performed with General Analysis 3 in Nikon Elements (5.20.00 or above). Briefly, a cell nuclei binary mask was created by Hoechst signal to calculate the number of cells in each field of view. CPSF6 localization and quantification was determined by creating binary masks of CPSF6 within the cells. Cytoplasmic CPSF6 was determined by subtracting the CPSF6 binaries from ones colocalized with Hoechst (nucleus) signal. Mean intensity and volume were recorded for each binary. Virus localization was determined by the Spot Detection function to create binary masks for spots positive for mRuby3/tagRFP, FAP-GPI, CypA-DsRed, or p24 signals. Trafficking data of HIV-1 was determined by using the Track Function with the spot binaries in random and constant motion mode. Any tracks with less than 20 frames were excluded in the data analysis.

### Protein expression and purification

The full length CPSF6 protein was expressed in a suspension-adapted HEK293 cell line (Expi 293F, Thermo Fisher Scientific) by transfection of expression plasmid using ExpiFectamine 293 (Thermo Fisher Scientific) according to the manufacturer’s instructions. Following transfection, the cells were grown at 37°C by shaking at 125 rpm in 8% CO2, 80% humidity for 2 days. The cells were harvested after 48 h of transfection by centrifugation at 100g for 10 min. The cell pellet was washed with cold PBS and flash-frozen and stored at −80° C.

The thawed cell pellet was resuspended in buffer A (50 mM HEPES-KOH (pH 8), 500 mM NaCl, and 5% glycerol, 2 mM DTT) supplemented with detergents (1% Tween 20 and 0.3% NP-40), deoxyribonuclease I (50 μg/ml; Sigma Aldrich) in the presence of a cocktail of protease inhibitors (Roche). After 2 h of rotation at 4° C, the lysate was homogenized by 15 strokes in an ice-cold, tight-fitting Dounce homogenizer. The homogenate was then centrifuged at 21,000*g* at 4°C for 30 min. After centrifugation, the supernatant was collected and mixed with 1 ml of amylose agarose resin (New England Biolabs) pre-equilibrated with buffer A per 50 ml of cell homogenate. The mixture was incubated with rotation at 4° C for 2 h and then transferred to a column. The resin was washed with 50x resin volume of buffer A. To elute the recombinant protein, the resin was incubated, in batch, with buffer A containing 100 mM maltose for 15 min at 4° C, and the flow through was collected as eluate. The eluate was applied to a Hi-Load Superdex 200 16/60 column (GE Healthcare) in a buffer A, and fractions containing target protein were collected and concentrated to 6-8 mg/ml using Amicon concentrators (Millipore, Billerica, MA, USA), flash-frozen with liquid nitrogen, and stored at −80° C.

His_6_-tagged CypA-DsRED-Exp2 was expressed in *E. coli* Rosetta 2 (DE3) cells (EMD Millipore) with autoinduction medium at 18° C for 16 h. Protein was purified using 5 mL Ni-NTA column (GE Healthcare) and a HiLoad Superdex200 16/60 size exclusion column (GE Healthcare) equilibrated with a buffer containing 25 mM sodium phosphate, pH 7.5, 150 mM NaCl, 1 mM dithiothreitol (DTT), 10% glycerol, and 0.02% sodium azide.

### SDS-PAGE and western blot analysis

For *in vitro* CPSF6 experiments, an equal volume of cell lysate and each fraction from the column were mixed with 4x NuPAGE LDS sample buffer (Thermo Fisher Scientific) supplemented with 10 mM DTT and loaded onto a 10% Bis-Tris NuPAGE gel (Thermo Fisher Scientific), alongside a protein molecular weight marker (BLUEstain protein ladder, Gold Biotechnology). Gels were run at 100 V for 15 min and then 150 V for 40 min in NuPAGE MES SDS running buffer and the proteins were subsequently transferred onto PVDF or nitrocellulose membranes using iBlot transfer stacks (Thermo Fisher Scientific). The membranes were blocked at ambient temperature for 1 h in BSA blocking buffers, followed by overnight at 4° C with rabbit anti-Maltose Binding Protein antibody (ab9084, Abcam) or rabbit anti-CPSF6 antibody (EPR12898, Abcam), then a further hour with monoclonal anti-rabbit immunoglobulins−alkaline phosphatase antibody at ambient temperature. Between each antibody incubation, the membranes were washed three times with TBS buffer containing 0.1% Tween 20 and finally the membranes were developed with BCIP/NBT color development substrate (Promega) to enable visualization of protein bands (Promega, USA). Each experiment was performed at least three times.

For measurement of CPSF6 expression in cells, an equal number of transduced and puromycin-selected HeLa cells or PBMCs were lysed with RIPA buffer (Bio-Rad), mixed with sample buffer (Bio-Rad), and to 100° C for 5 min. Denatured cell lysate was run on pre-casted 4-15% Criterion Tris-HCl gels (Bio-Rad) at 150 V for 1.5 h. Proteins were transferred to nitrocellulose membranes using a semi-dry transfer apparatus (Thermo Fisher Scientific) at 160 mA for 1 h. The membranes were blocked with 5% milk in PBS containing 0.1% Tween-20 at room temperature for 20 min. Primary antibodies anti-CPSF6 (NBP1-85676) and anti-α-tubulin (T5168, Sigma Aldrich) were used with secondary anti-mouse IgG or anti-rabbit IgG conjugated with horseradish peroxidase antibodies (A9917 and AP132P, Sigma Aldrich). SuperSignal West Pico Chemiluminescent substrate (Thermo Fisher Scientific) was used to visualize protein bands with Amersham Hyperfilm (GE).

### Capsid binding assay

Tubular assemblies of WT HIV-1 CA protein were prepared at 80 µM (2 mg/ml) in 1 M NaCl and 50 mM Tris-HCl (pH 8.0) buffer at 37° C for 1 h. N74D CA was dialyzed against 1 M NaCl and 50 mM Tris-HCl (pH 8.0) buffer at 4° C overnight at the concentration of 20 mg/ml. Before binding, the assembled mixture was diluted to 80 µM (2 mg/ml). For the binding assays, the binding buffer was the same as the stock buffer for MBP-CPSF6 proteins described above. Different concentrations of MBP-CPSF6 were added to preassembled CA tubes at CA concentration of 64 µM. The reaction mixtures were incubated on a rocking platform at room temperature for 1 h with gentle mixing at 10 min intervals. At the end of incubation, 5 µl samples were withdrawn from the reaction mixtures and immediately used for EM analysis. The remaining samples were pelleted at 21,000 g with a SORVAL Legend micro 21R centrifuge (Thermo Fisher Scientific) for 30 min and supernatants (s) and pellets (p, resuspended in the same volume) were mixed with 4x LDS loading buffer for gel analysis. Supernatant and pellet samples, without boiling, were loaded on 10% SDS-PAGE and stained with Coomassie Blue. Each experiment was performed at least three times.

To determine the binding ratio of MBP-CPSF6:CA, SDS-PAGE gels were scanned using an Epson 4990 scanner. The integrated intensities of CA and MBP-CPSF6 protein bands were measured using Image J 1.40 program (NIH). The molar ratios were calculated according to the formula (MBP-CPSF6 intensity/MBP-CPSF6 molecular weight)/(CA intensity/CA molecular weight) and calibrated using the input ratios as standards.

For binding of CypA and MBP-CPSF6 with HIV-1 capsid, 5 µM of CypA-DsRed was added to 10 µM pre-assembled WT CA-NC tubes, and at the same time 15 µM of competitive inhibitor CsA was added as a negative control. The reaction mixtures were incubated on a rocking platform at room temperature for 1 h with gentle mixing at 10 min intervals. 5 µM of MBP-CPSF6 P1 or P2 was added to the reaction and incubated on a rocking platform at room temperature for 1 h with gentle mixing at 10 min intervals. At the end of the incubation, the samples were pelleted as described above. Each experiment was performed at least three times. Nikon Elements 5.0 was used to quantify the binding ratio of CypA and MBP-CPSF6 with pre-assembled CA-NC tubes.

### TEM analysis

The morphologies of different variants of CA assemblies and CA–MBP-CPSF6 complexes were characterized by TEM. Samples were stained with fresh 2% uranyl formate, deposited onto 400-mesh carbon-coated copper grids, and dried for 30 min. TEM images were acquired on a Tecnai T12 transmission electron microscope at 120 kV.

### Type I IFN assay

To measure type I IFN from infected cells, HEK93-ISRE-luc cells were incubated with Cellstriper buffer (Thermo Fisher Scientific) for 30 min at room temperature to detach the cells and plated in 96-well plates. Media from infected cultures were added to cells in duplicate. Cells were lysed 21 h later and assessed for luciferase production. Dilutions of IFNα (Sigma-Aldrich) was included as a standard (0.001 – 2 U/ml).

### Statistics

Each virus infection experiment and associated imaging analysis was performed in at least two separate replicates. Compiled data was obtained from minimally two independent experiments or three donors. Statistical significance was determined by two-sided unpaired Student’s *t* test using Prism (GraphPad). *P* values of <0.05 were considered statistically significant. *P* values > 0.05, ns; 0.01 - 0.05, *; 0.01 - 0.001, **; 0.001 - 0.0001, ***; < 0.0001, ****.

## Acknowledgements

The authors thank Stephanie Ander, Douglas Fischer, Sharie Ganchua, and Chris Kline, for technical assistance; and Rahm Gummuluru, Vineet KewalRamani, Ned Landau, Greg Melikian, and Greg Towers for reagents. This work was supported by National Institutes of Health (NIH) P50 grant AI150481 (A.N.E., P.Z., S.C.W., and Z.A.), NIH R01 grant AI052014 (A.N.E.), NIH R01 grant GM114075 (C.T. and M.P.B.), NIH T32 training grant AI065380 (E.A.B.), and the UK Wellcome Trust Investigator Award 206422/Z/17/Z (P.Z.).

