## Supplemental figures for "Cytoplasmic CPSF6 regulates HIV-1 capsid trafficking and infection in a cyclophilin A-dependent manner"

**Supplemental Movie Legends**

**Movie S1. CPSF6-GFP traffics on microtubules.** Live-cell HILO imaging is shown of a CPSF6-GFP (green) higher-order complex trafficking on a microtubule stained with SiR (red) in a HeLa cell. (See also Figure S1.)

**Movie S2. HIV-1 complexes traffic on microtubules.** Live-cell HILO imaging of mRuby3-IN particles (red) with SiR-stained microtubules (white) in a HeLa cell after infection with WT HIV-1. Arrowheads point to 2 mRuby3-IN complexes trafficking on microtubules.

**Movie S3. HIV-1 complexes associate and traffic with CPSF6-GFP in the cytoplasm.** Live-cell HILO imaging from the bottom of a HeLa cell expressing CPSF6-GFP (green) and infected with WT HIV-1 containing mRuby3-IN (red).

**Movie S4. CPSF6-iRFPtraffics with GFP-TNPO3.** Live-cell HILO imaging is shown of a HeLa cell expressing CPSF6-iRFP (red) and GFP-TNPO3 (green).

**Movie S5. FAP-GPI on an HIV-1 particle is lost over time after infection of a HeLa cell.** Live-cell HILO imaging from the bottom of a HeLa cell expressing CPSF6-GFP (green) and infected with WT HIV-1 containing mRuby3-IN (red) and FAP-GPI (white) in the presence of a FAP dye. A double-labeled particle is seen in which the FAP-GPI signal separates from the mRuby3-IN signal, suggesting loss of the virus membrane, which contains FAP-GPI, after fusion with the endosome.

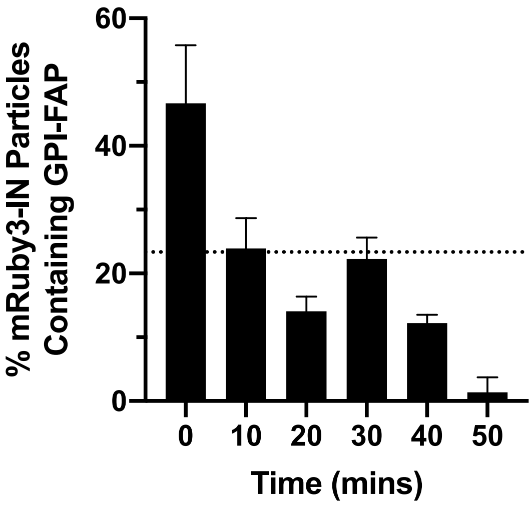


**Figure S1. FAP-GPI on HIV-1 particles is lost over time after incubation on HeLa cells.** HeLa cells were synchronously infected with WT HIV-1 produced with mRuby3-IN and FAP-GPI and imaged by live-cell microscopy (see Movie S1). The average percentage of mRuby3-IN complexes still containing FAP-GPI was determined between 0 - 50 min from n ≥ 75 cells/experiment from two separate experiments. Error bars represent STDEV and the dotted line represents 50% of double positive HIV-1 complexes at time 0.

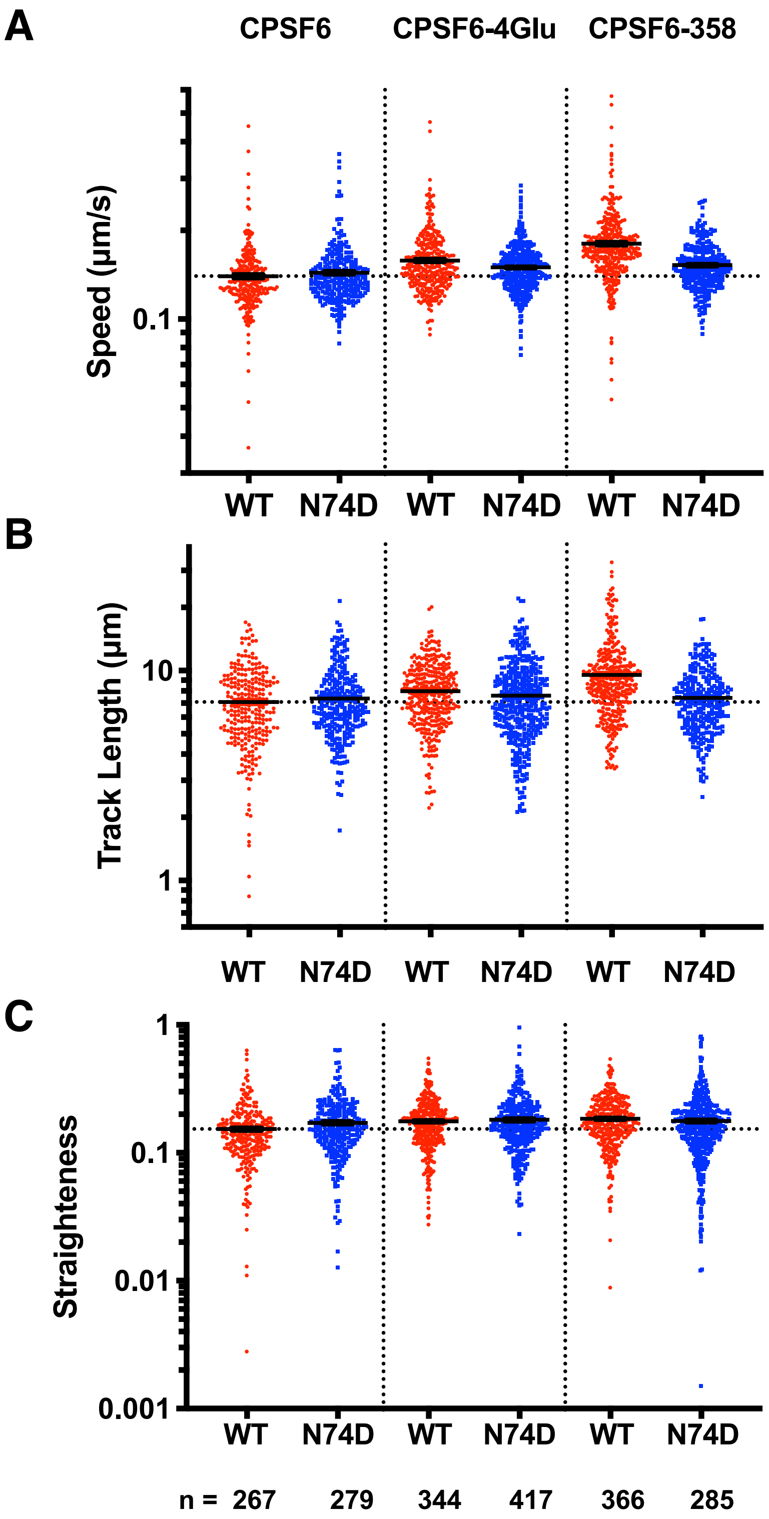


**Figure S2. Mutation or truncation of the CPSF6 R/S domain alters HIV-1 trafficking.** Results from HILO live-cell imaging are shown that were summarized in Figure 3D. The particle speed (A), track length (B), and track straightness (C) of individual WT or N74D HIV-1 mRuby3-IN complexes in HeLa cells expressing CPSF6-iRFP, CPSF6-4Glu-iRFP, or CPSF6-358-iRFP are shown. Error bars indicate SEM. Dotted lines denote the mean of WT complexes in CPSF6-iRFP cells. The number (n) of mRuby3-IN complexes analyzed for each condition are listed at the bottom.

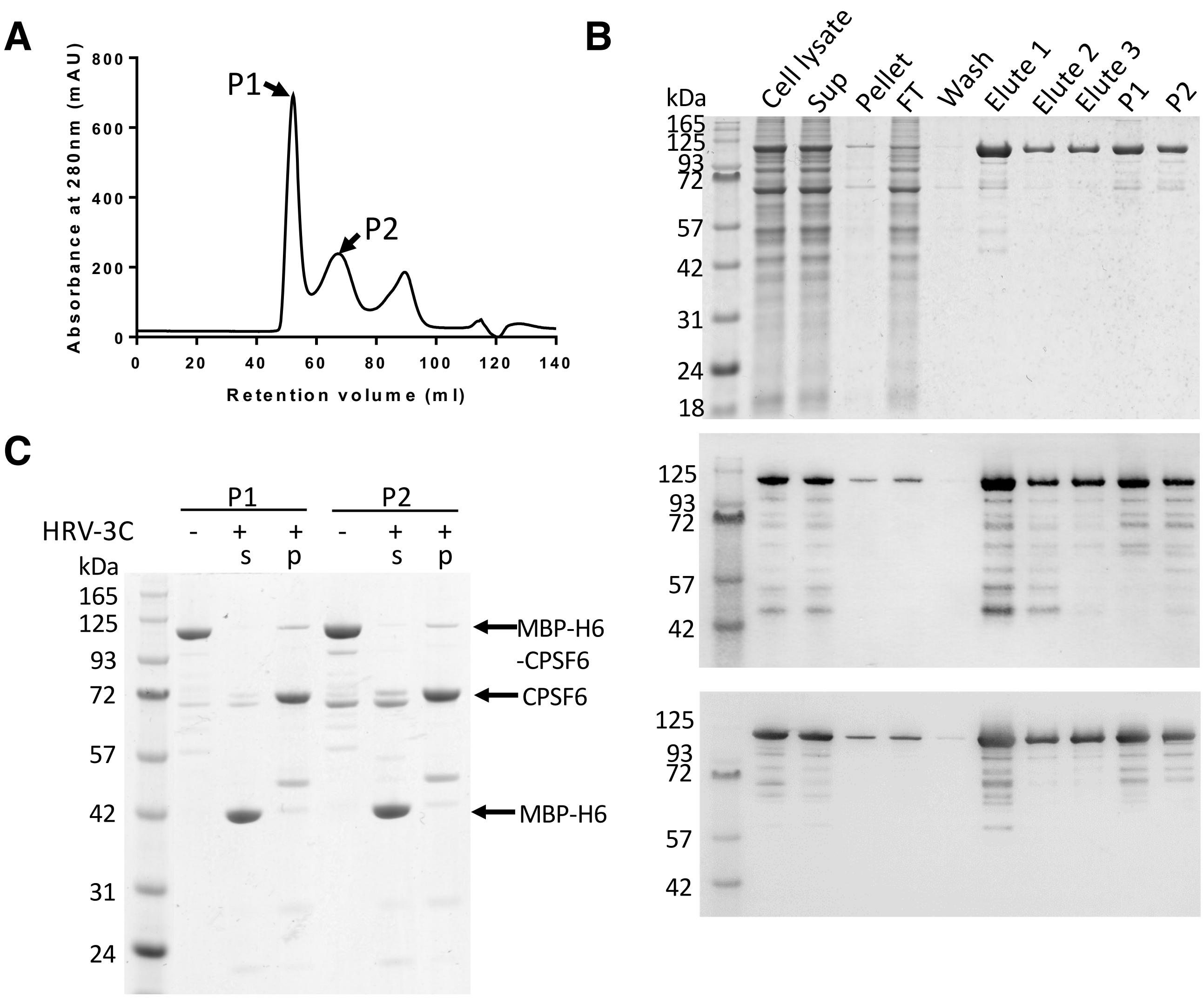


**Figure S3. Purification of MBP-CPSF6 with an MBP tag from mammalian cell expression system.** (A) Gel filtration profile of the protein eluted from the Superdex 200 16/60 column. The two MBP-CPSF6 peaks are labeled P1 and P2. (B) SDS-PAGE and Western blot analysis of MBP-CPSF6 purification. Samples were taken from cell lysate, supernatant (sup), pellet, flowthrough (FT), wash, elute 1, 2, and 3 from Amylose resin, and peaks (P1 and P2) from the Superdex 200 16/60 column (showed in panel A) were stained with Coomassie blue (top) or processed with anti-MBP (middle) or anti-CPSF6 (bottom) antibody, following Western blotting. (C) MBP tag removal analysis, the uncleaved P1 and P2 was shown in lane 2 and 5, the supernatant (s) and pellet (p) of P1 and P2 after cleavage with HRV-3C protease were shown in lane 3, 4, 6, and 7, respectively. Samples were stained with Coomassie blue. Proteins are indicated by arrows on the right.

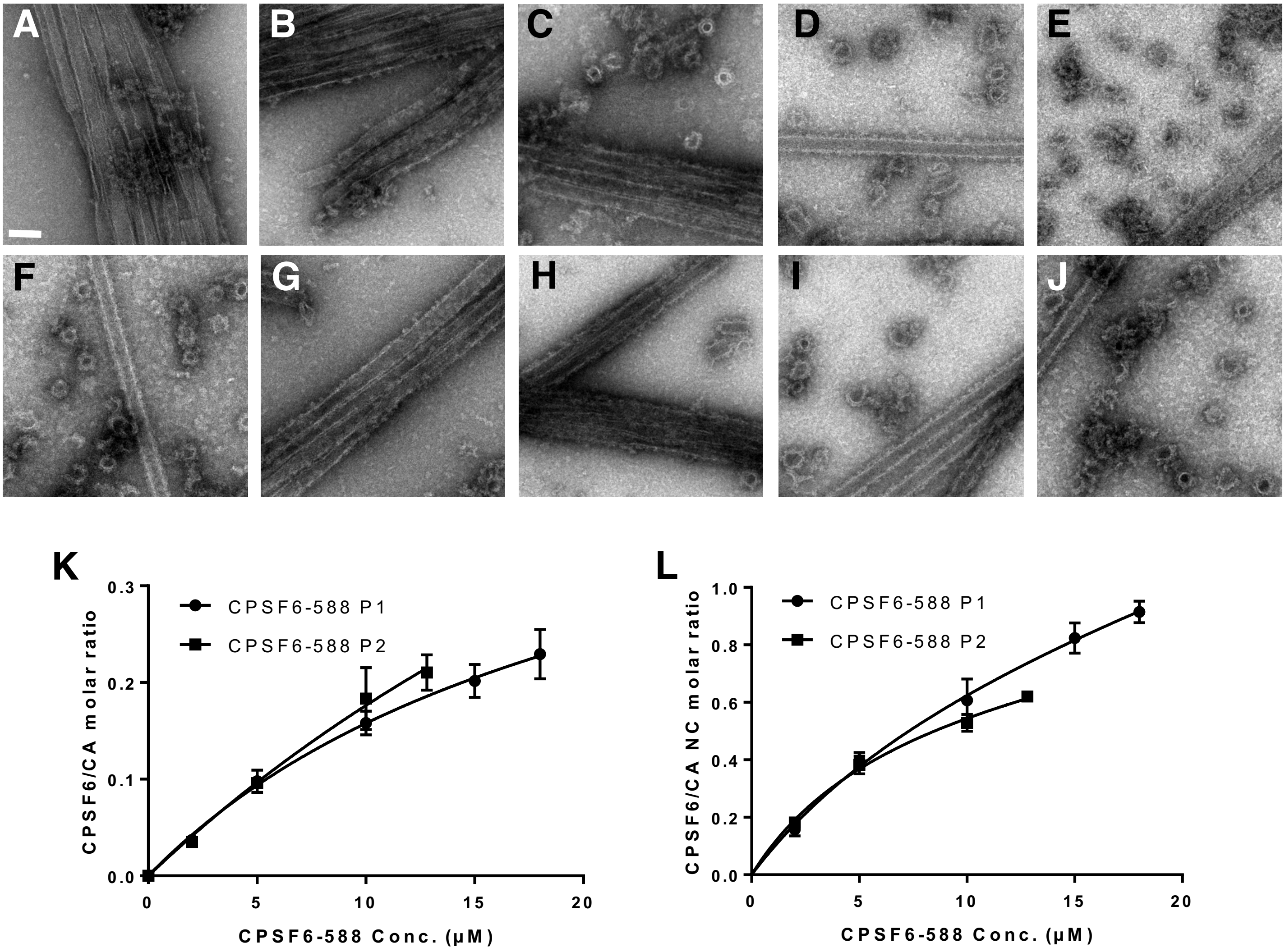


**Figure S4. Dose dependent effect of MBP-CPSF6 on HIV-1 capsid tubes.** (A-J) Representative negative-stain EM micrographs of CA with different concentrations of MBP-CPSF6 P1 or P2. WT CA tubular assemblies alone (A) or with 2 μM (B), 5 μM (C), 10 μM (D), 15 μM (E), or 18 μM (F) of MBP-CPSF6 P1, or with 2 μM (G), 5 μM (H), 10 μM (I), or 12.8 μM (J) of MBP-CPSF6 P2. Scale bars, 100nm. (K&L) Dose dependent effect of MBP-CPSF6 on CA tubes (K) or CA-NC tubes (L). Shown is binding of P1 (circles) and P2 (squares) to assembled WT CA tubes or CA-NC tubes. The error bars indicate the standard deviation of the values.

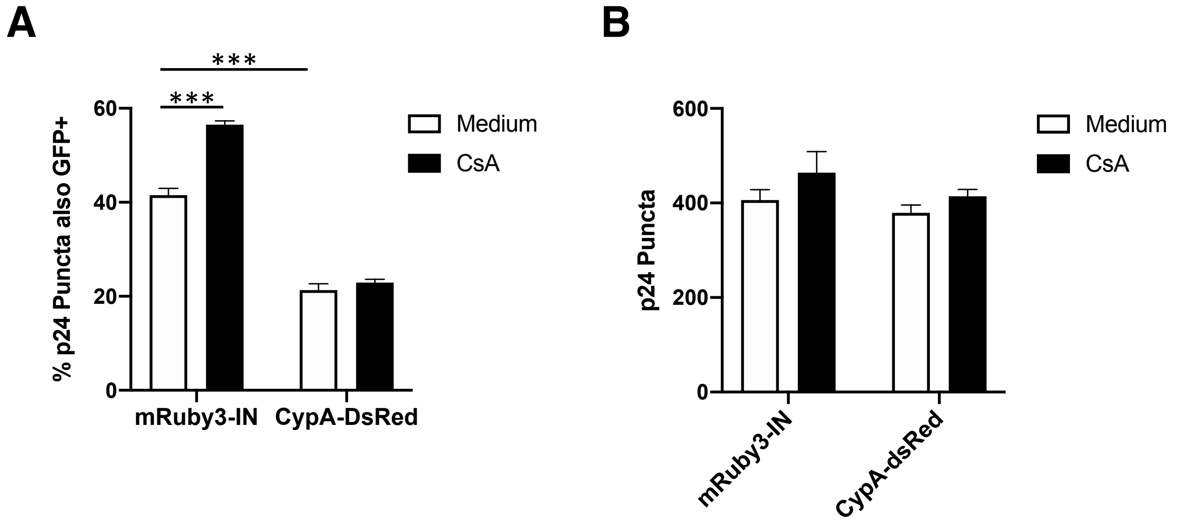


**Figure S5. CypA-DsRed prevents formation of HIV-1-induced CPSF6-358-GFP higher order complexes.** HeLa cells were infected with WT HIV-1 containing mRuby-IN or CypA-DsRed in the presence or absence of 10 µM CsA for 1 h. Cells were fixed and stained with p24 antibodies. (A) The percentage red viral complexes that were also positive for GFP was plotted. (B) The total number of p24+ particles in cells was plotted. Error bars represent STDEV.

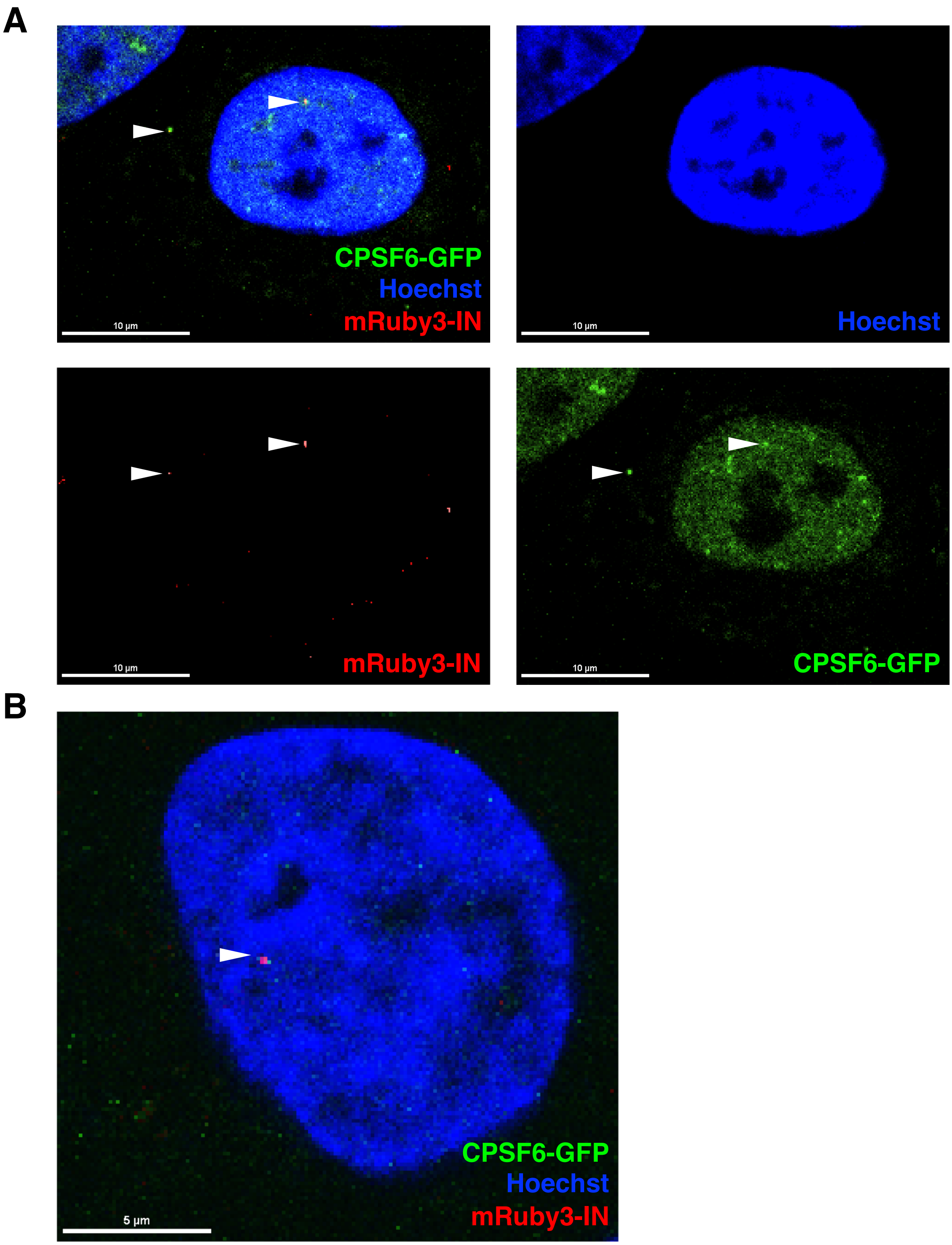


**Figure S6. WT HIV-1 induces formation of CPSF6-GFP puncta in nuclei.** HeLa cells expressing CPSF6-GFP were infected with WT HIV-1 containing mRuby3-IN and fixed after 2 h. (A) A representative cell is shown in which a mRuby3-IN complex is co-localized with CPSF6-GFP in the cytoplasm and another in the nucleus. (B) Another representative cell in which mRuby3-IN is co-localized with CPSF6-GFP in the nucleus. White bars, 5 µm.

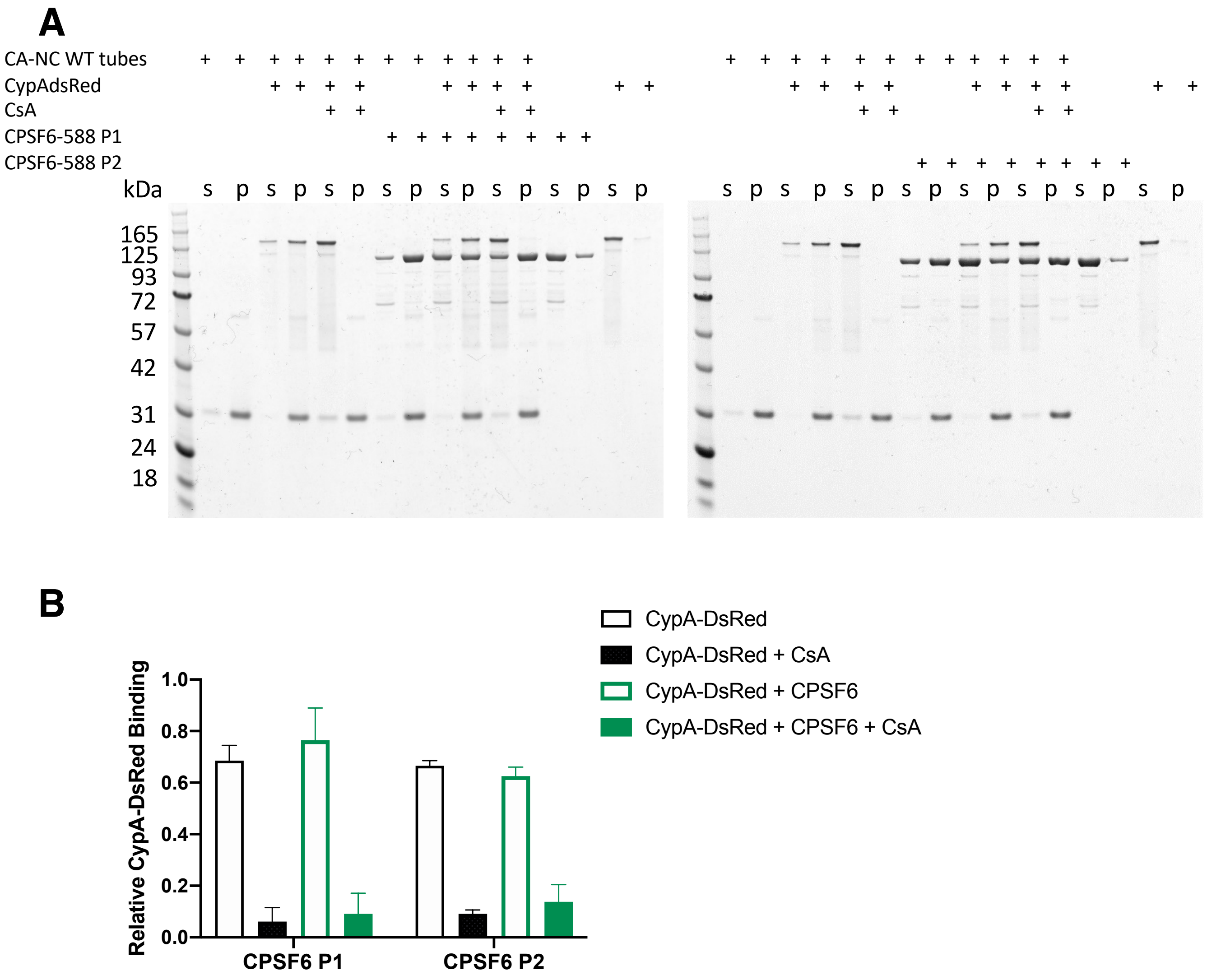


**Figure S7. CypA prevents HIV-1 capsid binding to CPSF6.** (A) SDS-PAGE of WT CA-NC assemblies were incubated with CypA-DsRed with or without CsA for 1hr at room temperature, following incubation with MBP-CPSF6 P1 or P2 and then centrifuged. The gel was Coomassie blue stained, with supernatant (s) and pellet (p) sample indicated. (B) The binding ratio of CypA-DsRed with WT CA-NC assemblies. The error bars indicate the standard deviation of the values of three independent experiments.

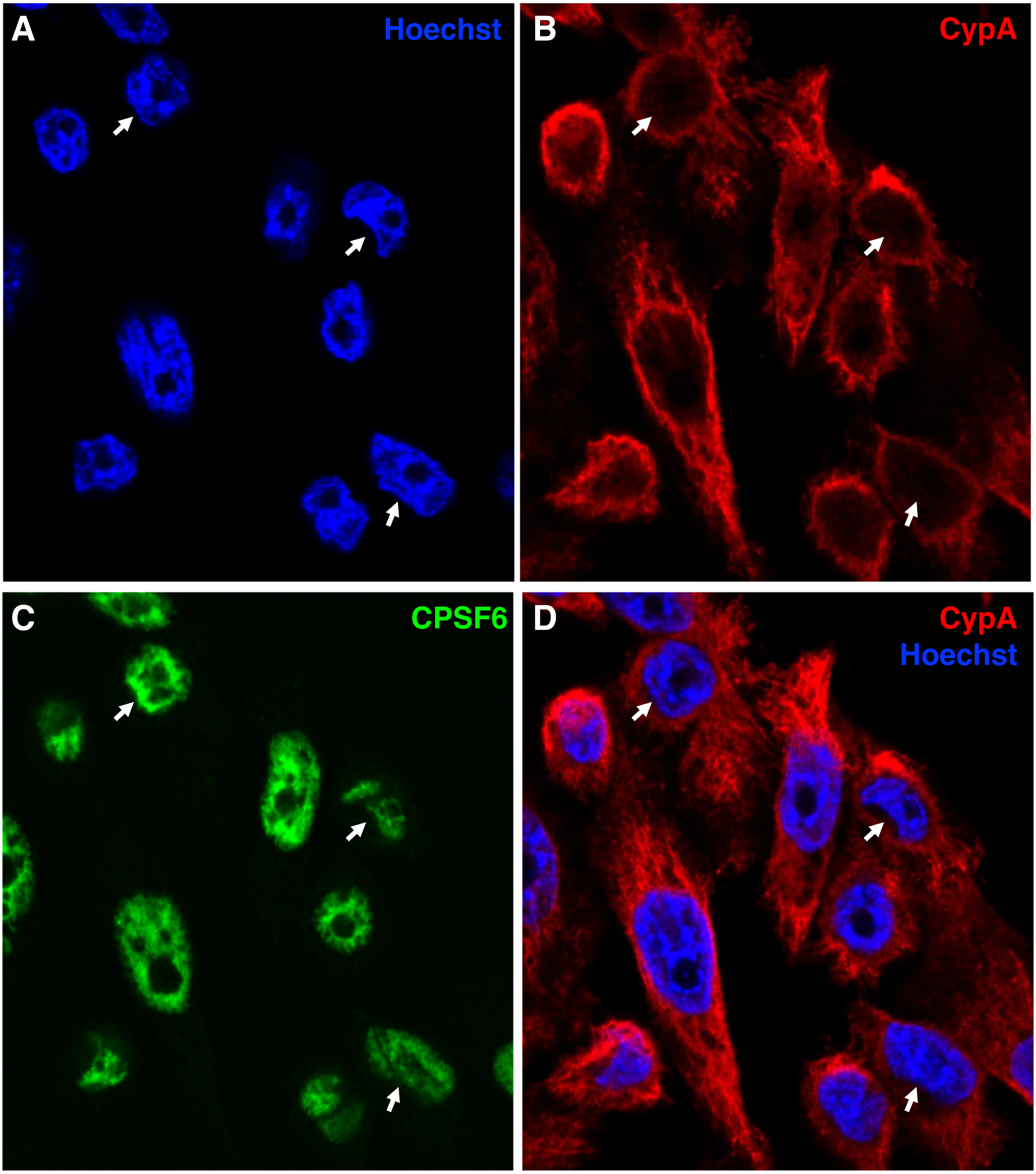


**Figure S8. CypA expression is excluded from the nucleus of HeLa cells.** Confocal micrographs are shown of HeLa cells stained with (A) Hoechst, (B) CypA antibodies, (C) CPSF6 antibodies, and (D) Hoechst and CypA antibodies. Arrows show perinuclear exclusion of CypA.

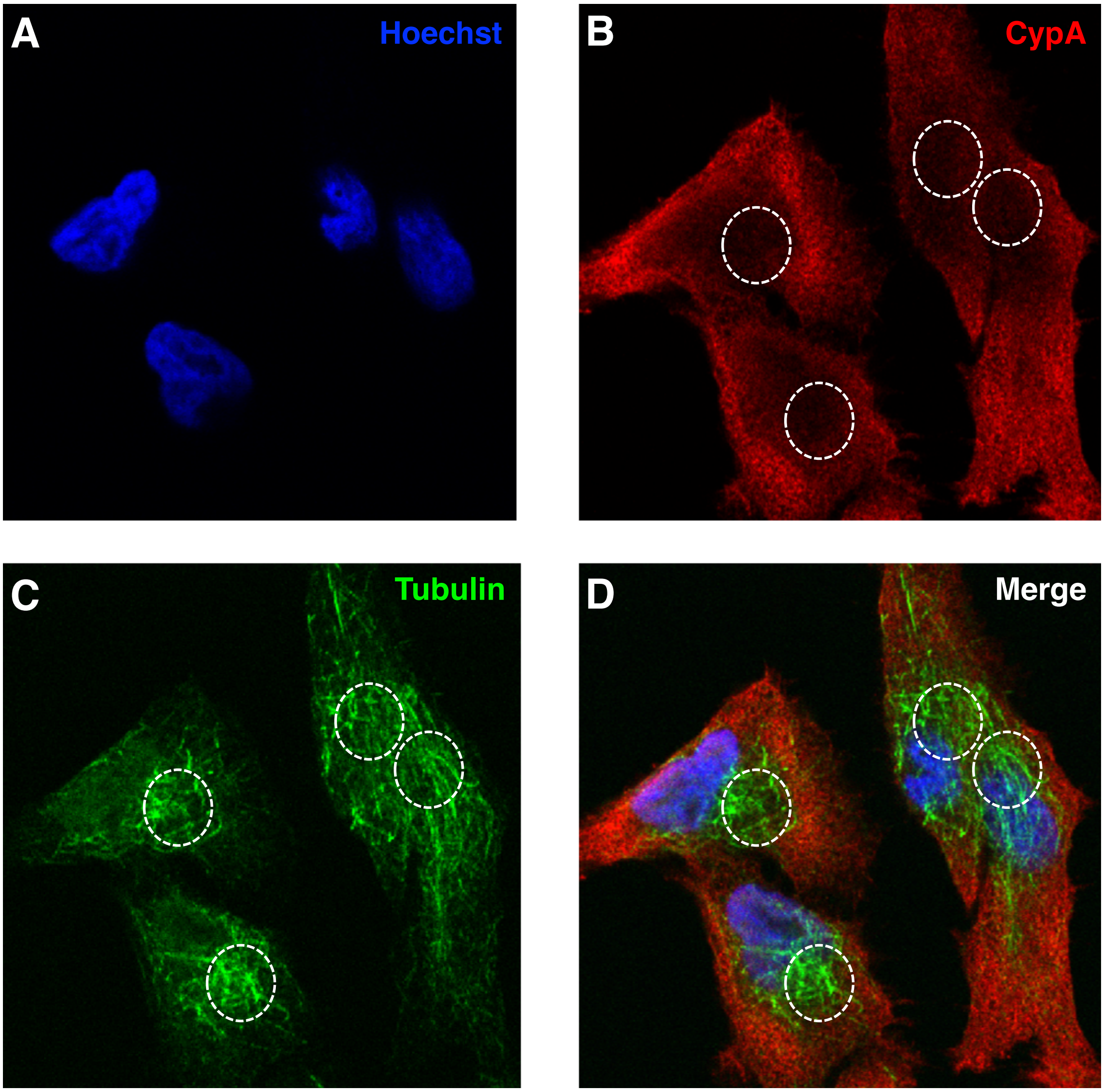


**Figure S9. CypA expression is excluded from the microtubule-organizing center of HeLa cells.** Confocal micrographs are shown of HeLa cells stained with (A) Hoechst, (B) CypA antibodies, (C) tubulin antibodies, and (D) all stains.

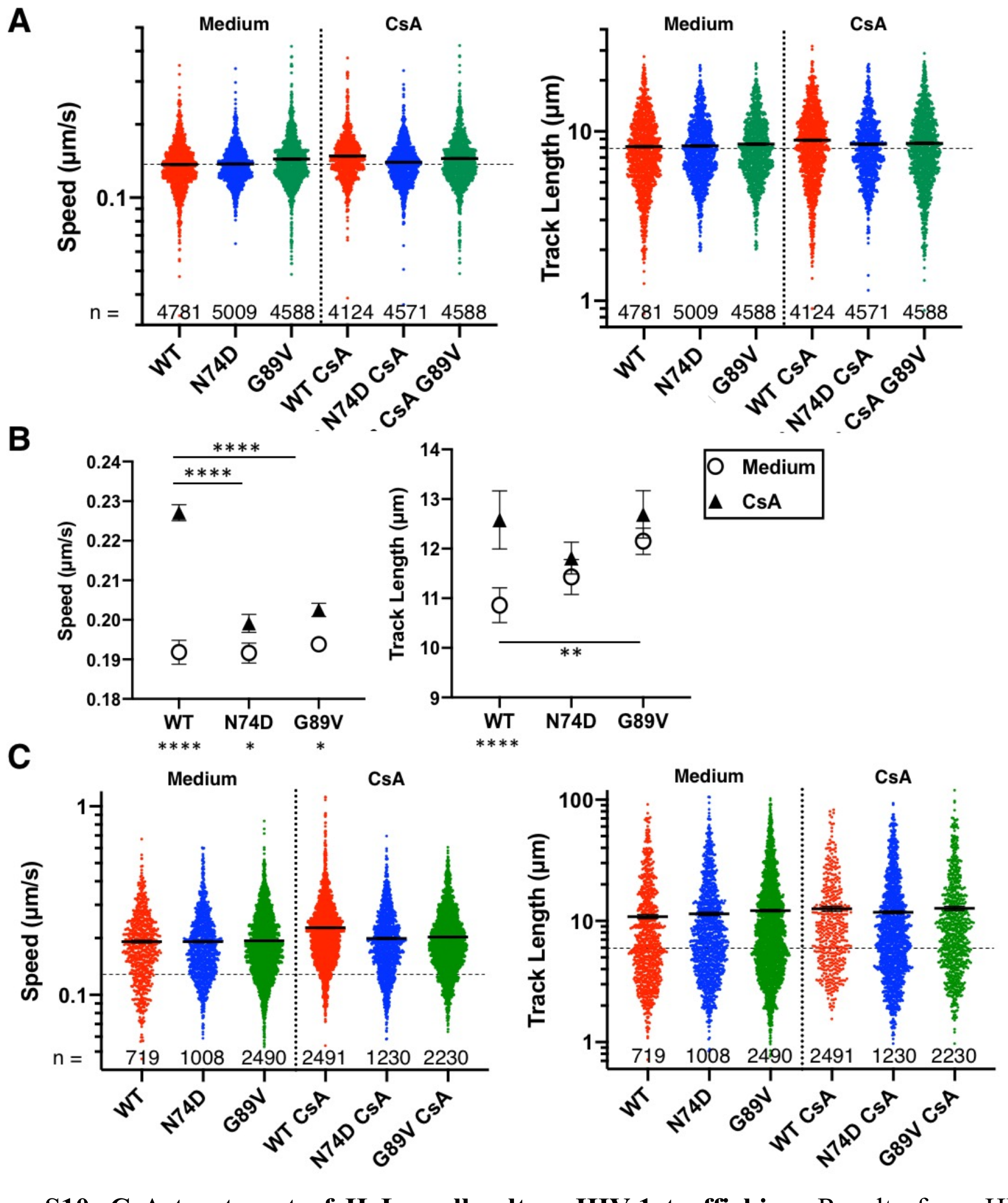


**Figure S10. CsA treatment of HeLa cells alters HIV-1 trafficking.** Results from HILO live-cell imaging are shown. (A) As summarized in Figure 5F, the speed and track length of individual WT, N74D, or G89V HIV-1 mRuby3-IN complexes in HeLa cells expressing treated with or without 10 µm CsA are shown. Dotted lines denote the mean of WT complexes. Error bars indicate SEM. The number (n) of mRuby3-IN complexes analyzed for each condition are listed at the bottom of the graph. (B) The average particle speeds and track lengths of WT, N74D, or G89V HIV-1 mRuby3-IN complexes in HeLa cells expressing CPSF6-GFP treated with or without 10 µm CsA are shown. Error bars indicate SEM. (C) The speed and track length of individual WT, N74D, or G89V HIV-1 mRuby3-IN complexes from (B) are shown. Dotted lines denote the mean of WT complexes. Error bars indicate SEM. The number (n) of mRuby3-IN complexes analyzed for each condition are listed at the bottom of the graph.

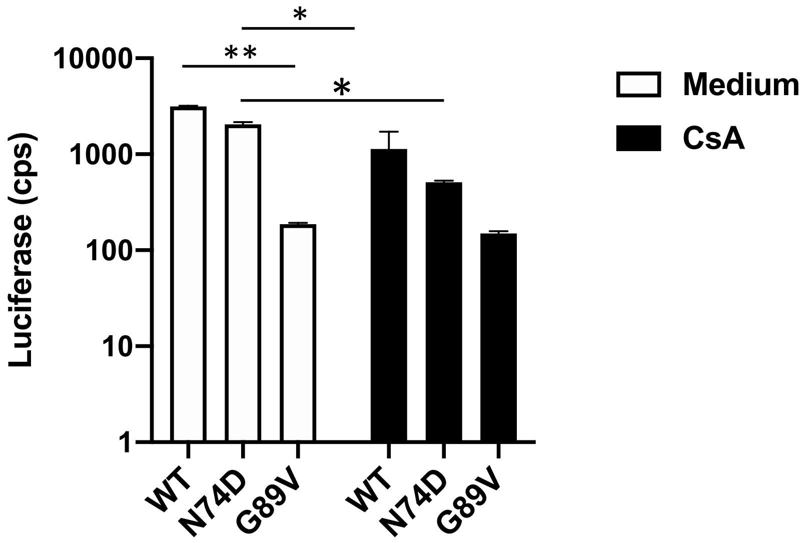


**Figure S11. CsA treatment of HeLa cells does not affect WT or G89V HIV-1 infection but does affect N74D HIV-1 infection.** HeLa cells were treated with 5 mM CsA and infected with WT, N74D, or G89V HIV-1. Results are shown as the mean luciferase expression of duplicates with error bars indicating STDEV.

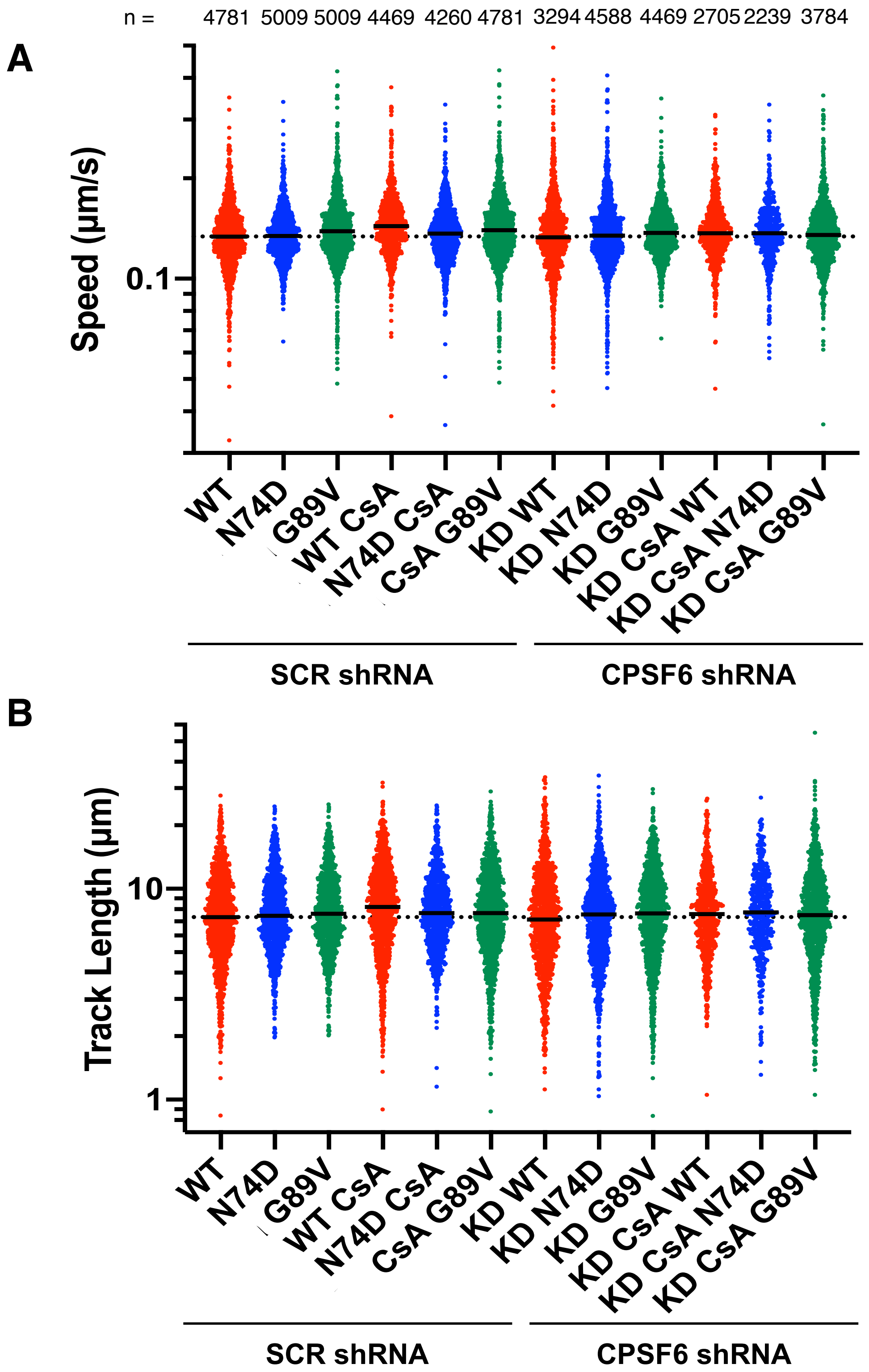


**Figure S12. CPSF6 knockdown in HeLa cells alters HIV-1 trafficking during CsA treatment.** Results from HILO live-cell imaging are shown that were summarized in Figure 6B. The particle speed (A) and track length (B) of individual WT, N74D, and G89V HIV-1 mRuby3-IN complexes in HeLa cells expressing CPSF6-GFP are shown. Prior to infection, cells were transduced with lentivirus expressing SCR or CPSF6 shRNA and treated with or without CsA. Error bars indicate standard errors of the mean (SEM). Dotted lines denote the mean of WT complexes in CPSF6-GFP cells. The number (n) of mRuby3-IN complexes analyzed for each condition are listed at the top.

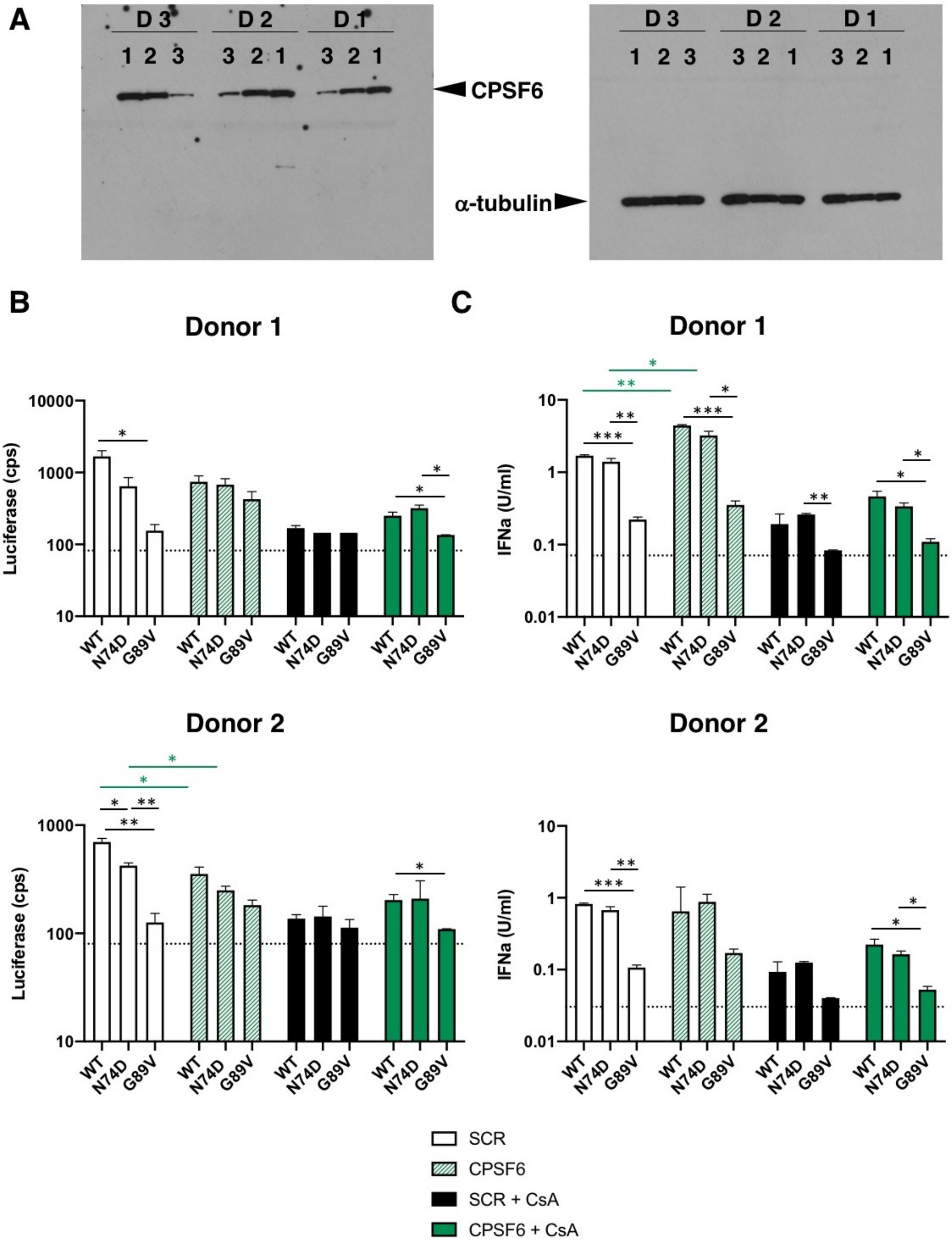


**Figure S13. CPSF6 knockdown in primary PBMC.** (A) CPSF6 depletion by shRNA was visualized in stimulated PBMCs by western blot of stimulated PBMCs. A scrambled shRNA was used as a knockdown control and α-tubulin was used as a loading control for the gel. Lane 1, no shRNA; Lane 2, scrambled shRNA; Lane 3, CPSF6 shRNA. (B) Infections with WT, N74D, and G89V HIV-1 were measured in primary PBMC cells from 2 donors after shRNA depletion of CPSF6, treatment with CsA, or both. (F) IFNα production from infected PBMC shown in (B) was quantified using HEK 293T ISRE-luc indicator cells.

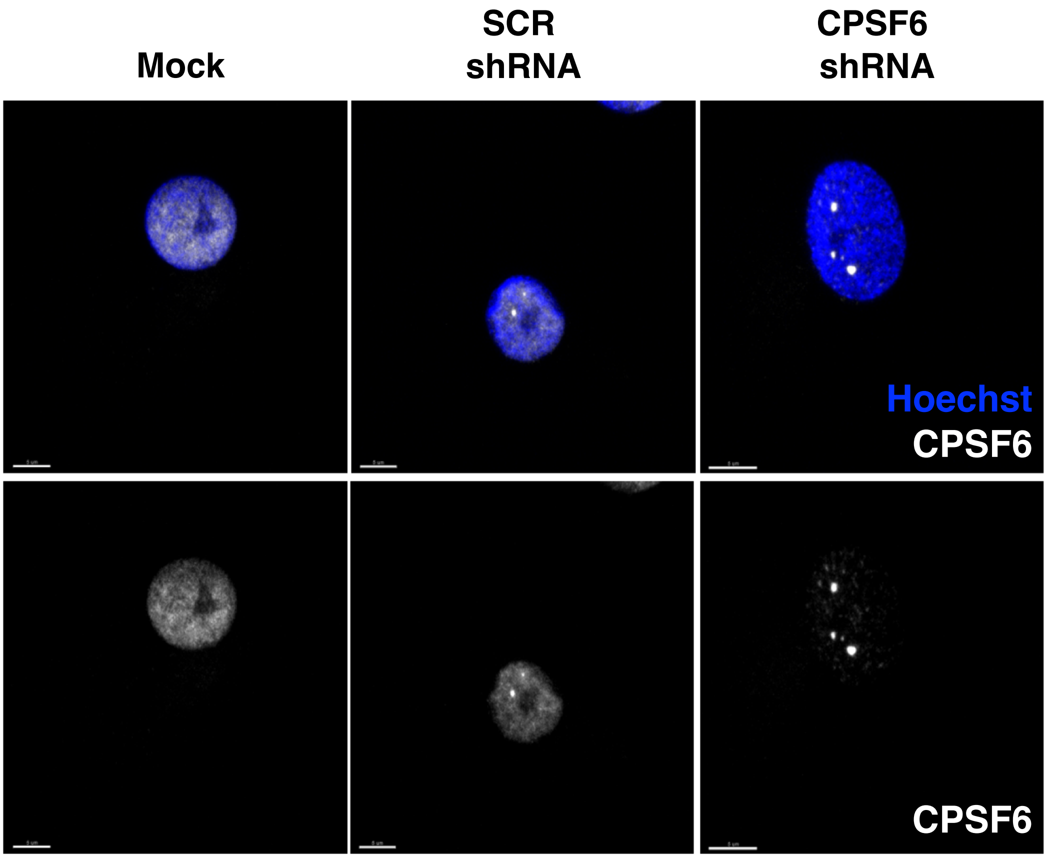


**Figure S14. Knockdown of CPSF6 in MDM via lentiviral shRNA transduction.** Representative confocal micrographs are shown for CPSF6 antibody staining in MDM without lentiviral transduction or transduction with lentiviruses expressing SCR shRNA or CPSF6 shRNA.

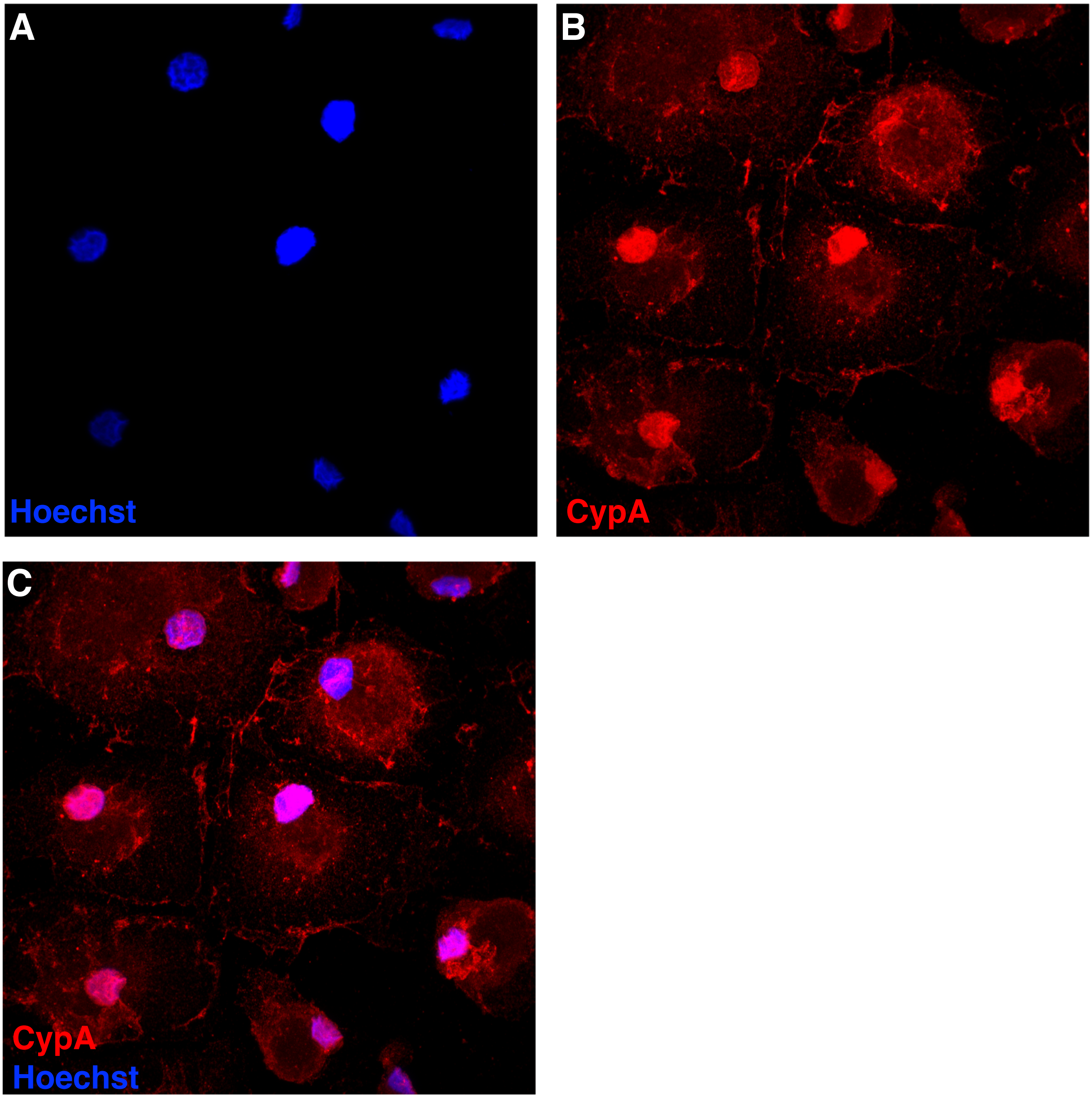


**Figure S15. CypA is expressed in the nucleus and cytoplasm of MDM.** Confocal micrographs are shown of MDM stained with (A) Hoechst, (B) CypA antibodies, (C) Hoechst and CypA antibodies.

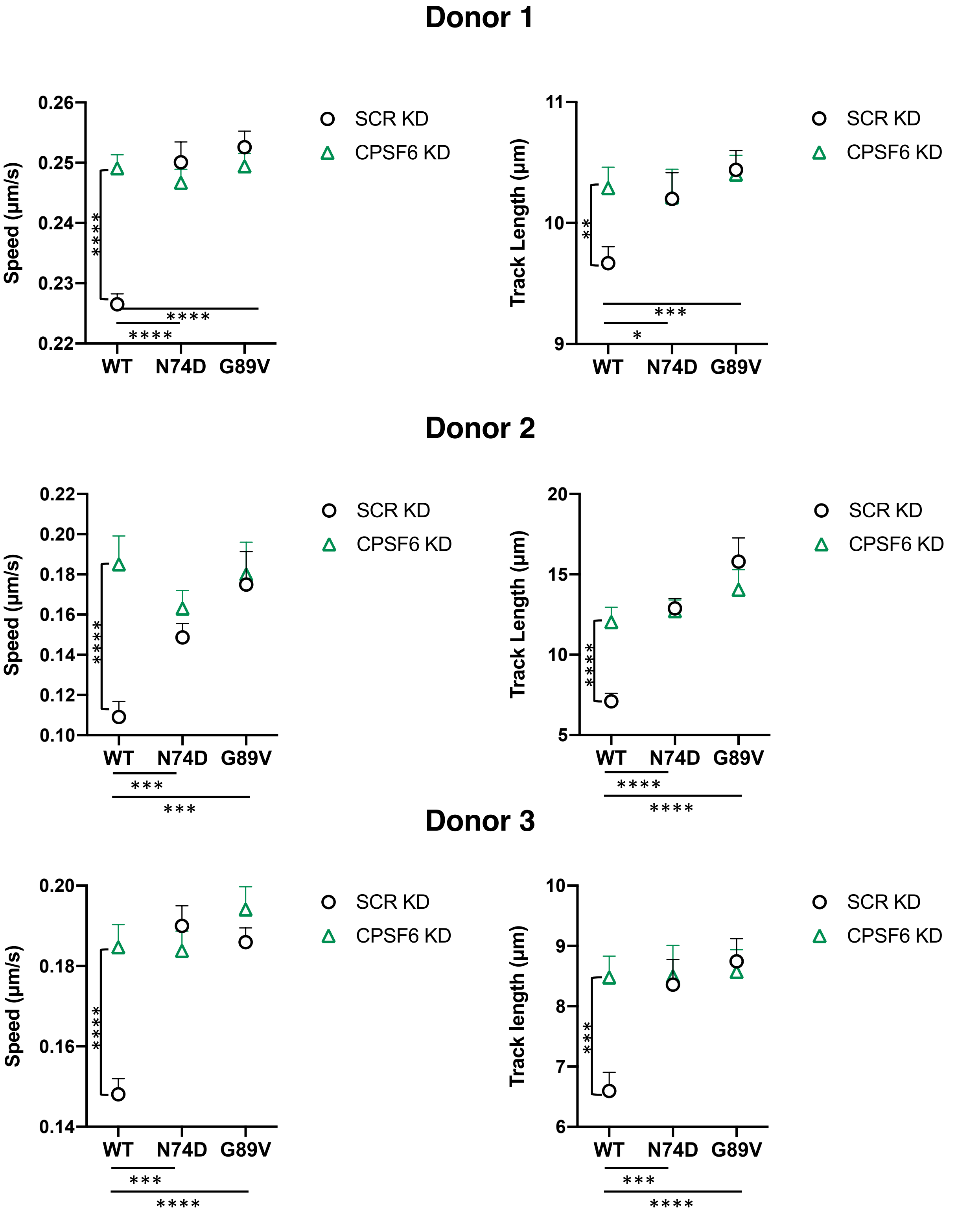


**Figure S16. CPSF6 knockdown alters HIV-1 trafficking in MDM.** The average particle speeds and track lengths of WT, N74D, or G89V HIV-1 mRuby3-IN complexes in MDM from 3 donors with SCR KD or CPSF6 KD are shown. Error bars indicate SEM.

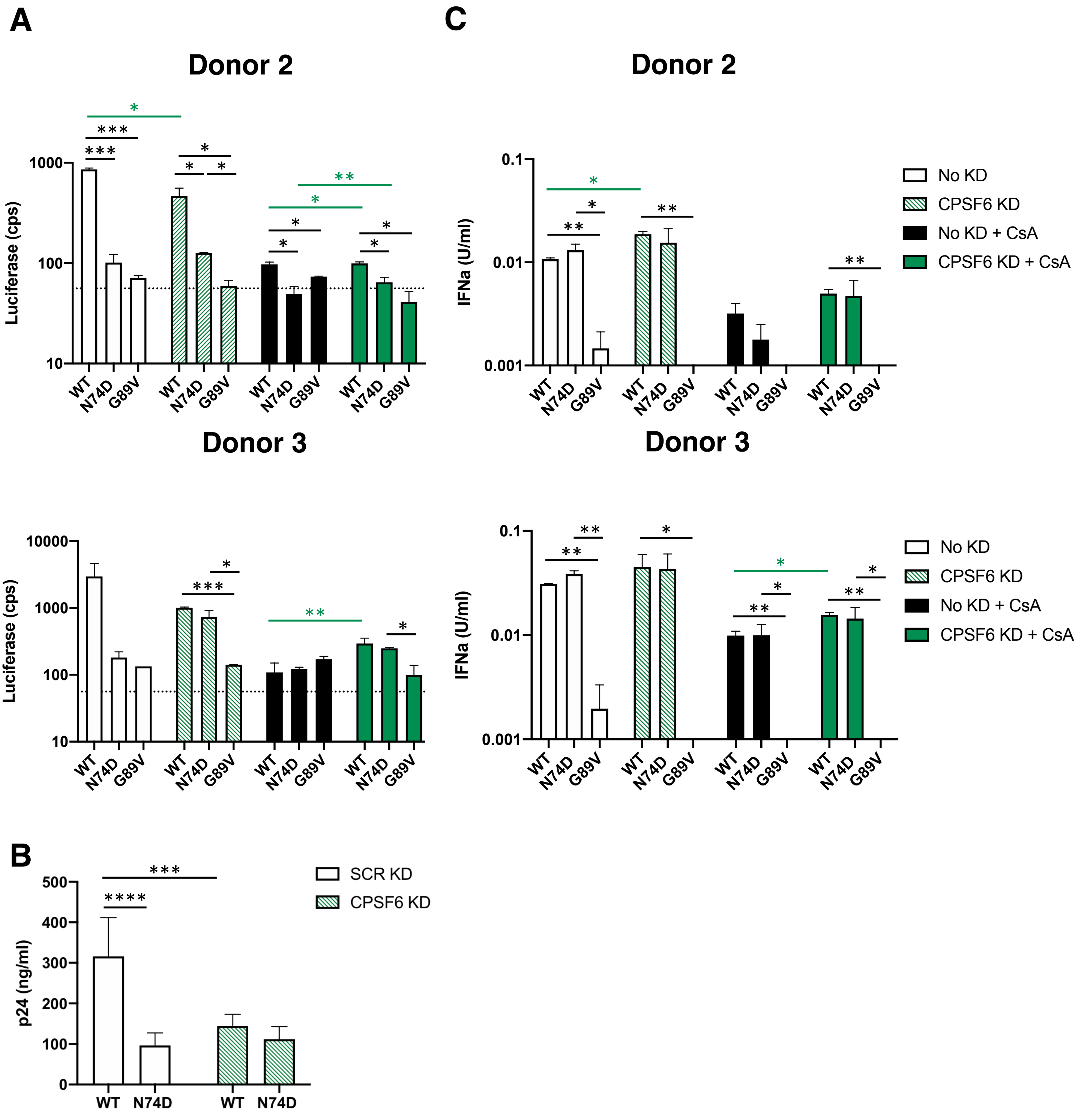


**Figure S17. CPSF6 knockdown or loss of CypA binding decreases HIV-1 infectivity in MDM.** (A) Infection of WT, N74D, or G89V HIV-1 in cells from Donor 2 and 3 shown in Figure S14 are graphed. Error bars indicate STDEV of duplicates and the dotted line represents the average luciferase expression of uninfected cells. (B) WT and N74D HIV-1$_{NL4\text{-}BAL}$ virus production of duplicate MDM infections was measured by p24 ELISA on d8 post-infection. Results are shown as means +/- STDEV. (C) IFNα was measured in the supernatants of the cells from (A). Error bars indicate STDEV of duplicates. The limit of detection is 0.001 U/ml.
